# Dendritic coincidence detection in Purkinje neurons of awake mice

**DOI:** 10.1101/2020.06.15.152496

**Authors:** Christopher J. Roome, Bernd Kuhn

## Abstract

Dendritic coincidence detection is thought fundamental to neuronal processing, yet the underlying dendritic voltage-calcium relationship remains unexplored in awake animals. Here, using simultaneous voltage and calcium two-photon imaging of Purkinje neuron spiny dendrites, we show how coincident sub- and suprathreshold synaptic inputs modulate dendritic calcium signaling during sensory stimulation in awake mice. Sensory stimulation evokes subthreshold excitatory and inhibitory post-synaptic potentials, that coincide with suprathreshold dendritic spikes triggered by climbing fiber and parallel fiber synaptic input. Purkinje neuron dendrites integrate these inputs in a time-dependent and non-linear fashion to enhance the sensory evoked dendritic calcium signal. Intrinsic supra-linear dendritic mechanisms, including voltage gated calcium channels and metabotropic glutamate receptors, are recruited cooperatively to expand the dynamic range of sensory evoked dendritic calcium signals. This establishes how dendrites use multiple interplaying mechanisms to perform coincidence detection, as a fundamental and ongoing feature of dendritic integration during behavior.

## Introduction

Dendritic integration is fundamental to signal processing in the brain. So far, most studies on dendritic integration were performed in vitro, in the absence of physiological inputs (Larkum et al., 2009; Markram et al., 1997; Stuart and Häusser, 2001; Wang et al., 2000). As such, our understanding of the basic components of dendritic integration; the frequency, amplitude, and spatio-temporal distribution of synaptic inputs under physiological conditions, and how these inputs are integrated by dendrites in awake behaving animals, remains incomplete.

Coincidence detection is a basic form of dendritic integration. By detecting coincident synaptic input, it is thought that neurons distinguish important signals from ongoing synaptic activity and modify synaptic strength through synaptic plasticity (Brown et al., 1990).

Purkinje neuron (PN) dendrites in the cerebellum are ideally suited to perform coincidence detection. They receive excitatory synaptic input from two distinct pathways; the climbing fiber (CF) and numerous parallel fibers (PF). CFs project from the inferior olive and evoke suprathreshold dendritic calcium spikes. PFs relay mossy fiber activity originating from the lateral reticular nucleus and pontine nuclei and together with synaptic input from inhibitory molecular layer interneurons (MLIs), evoke subthreshold postsynaptic potentials in the PN dendrites (Konnerth et al., 1990; Roome and Kuhn, 2018). In cerebellar slices, paired stimulation of these sub- and suprathreshold inputs evoke ‘supra-linear’ dendritic calcium signals in PNs, whereby the signal amplitude is larger than the sum of calcium signals triggered by PF and CF input alone. At PF-PN synapses supra-linear calcium signals lead to a form of synaptic plasticity, known as long-term depression (LTD) (Miyata et al., 2000; Wang et al., 2000).

This particular coincident detection process is well studied in PN spines and dendrites in cerebellar slices, and supports early theoretical predictions for motor learning in the cerebellum (Marr, 1969), but remains a controversial component in the theory of cerebellar function and motor learning (Mauk et al., 1998; Najafi and Medina, 2013; Sakurai, 1987; Schonewille et al., 2011; Wang et al., 2000). A corresponding detailed description of dendritic coincidence detection from awake animals is still missing, and the behavioral conditions under which these processes occur are unknown. Critically, coincidence detection of PF and CF input has not been confirmed in vivo, and recent studies have failed to detect supra-linear dendritic calcium signals triggered by coincident PF and CF input (Gaffield et al., 2019; Gaffield et al., 2018).

Due to technical limitations, PF evoked input to PN dendrites - that are predominantly subthreshold voltage signals - have been thus far undetectable in vivo. As such, confirming physiological interaction between PF and CF input has not been possible, and previous in vivo studies have instead focused on coincident MLI and CF activity and suprathreshold calcium signals only (Callaway et al., 1995; Gaffield et al., 2018; Kitamura and Häusser, 2011). We recently developed a technique to record both sub- and suprathreshold dendritic voltage signals, evoked by synaptic input in awake mice (Roome and Kuhn, 2018). Here by combining simultaneous voltage and calcium imaging from the PN spiny dendrites, we investigate sensory evoked dendritic signal processing and coincidence detection in PN dendrites awake mice.

## Results

Single PNs in lobule V of the cerebellar vermis were double labelled with voltage sensitive dye ANNINE-6plus and genetically encoded calcium indicator GCaMP6f, as described previously (Roome and Kuhn, 2018) (see Supplementary Information). Briefly, a chronic cranial window with access port (Roome and Kuhn, 2014) permitted two photon imaging, single neuron labelling and drug delivery (Figure 1a-c). In total 54 PNs in 23 mice were labelled and several PNs (up to 5) were labeled and recorded from in the same mouse (Figure 1b). Two photon linescan imaging (2kHz) recorded dendritic voltage and calcium signals simultaneously from the spiny PN dendrites, predominantly comprising PF-PN synapses, at a depth of 30-70μm below the dura (Figure 1c). Simultaneous voltage and calcium recordings were spatially averaged (across many dendritic processes) to maximize the signal to noise ratio, at a temporal resolution of 0.5ms/line. As the voltage response of ANNINE-6plus is linear down to a nanosecond time scale, the averaged voltage imaging signal represents the average membrane potential of dendritic shafts and dendritic spines within the point spread function of excitation along the scan line.

**Figure 1.**
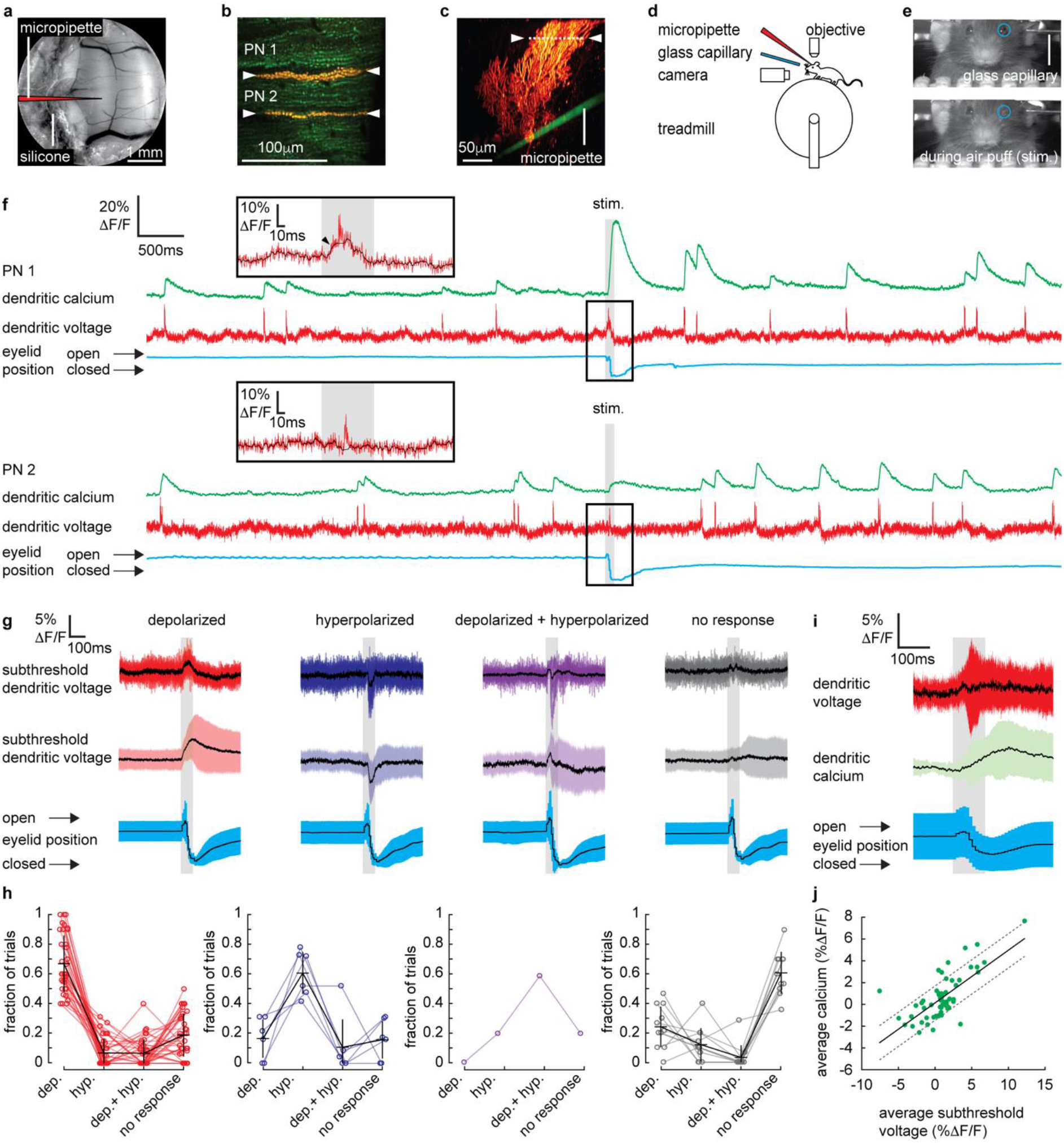
Coincident parallel fiber and climbing fiber synaptic input to Purkinje neuron dendrites during sensory evoked responses. (a) A chronic cranial window with access port on the cerebellar vermis lobule V was used for imaging from the dendrites of single PNs and allowed access to the brain via micropipette (schematically indicated). (b) 2P image of the spiny dendrites of two PNs (PN1 and PN2) labelled with voltage sensitive dye ANNINE-6plus (red) and genetically encoded calcium indicator GCaMP6f (green), resulting in double labelling (yellow). (c) Reconstruction of a single labelled PN showing position of 2P linescan (white dashed line and arrows) and the micropipette (green) positioned in the granular layer (50μm below the soma) used for pharmacological manipulation. (d) Sketch of the setup with a mouse mounted on a treadmill under a 2P microscope for imaging in lobule V of the cerebellar vermis. A glass capillary was used to deliver a 100ms air puff directed towards the ipsilateral eye. (e) A camera was used to monitor mouse movements and record eye responses during the air puff. Blue circles show ROIs used to record eyelid movement. (f) Single trial recordings of simultaneous dendritic voltage (red traces) and calcium (green traces) from PN1 and PN2 shown in (c). Suprathreshold signals are clearly visible in both voltage and calcium traces. Blue traces show average intensity of ROI used to record eye responses. Grey bars show full duration of air puff stimulus. The calcium recording from PN1 shows a dramatically enhanced evoked calcium signal during the air puff while the calcium recording from PN2 does not. Insets show zoomed voltage traces during air puff stimulation. The voltage recording in PN1 inset shows depolarizing subthreshold voltage signal, during the air puff (small black arrow and black trace; 10ms boxcar filtered voltage recording), while the voltage recording from PN2 shows no subthreshold signal. (g) Subthreshold voltage signals during air puff sorted into ‘depolarized’ (red), ‘hyperpolarized’ (blue), ‘depolarized + hyperpolarized’ (purple) and ‘no response’ (grey) groups, showing single recordings from a single PN (top traces; black trace shows mean) and averages all recordings in each group (middle traces; black trace shows mean, colored trace shows SD) and corresponding eyelid responses (bottom traces; black trace shows mean, colored trace shows SD). (h) Fraction of recordings resulting in each response type; depolarized (‘dep.’), hyperpolarized (‘hyp.’), depolarized + hyperpolarized (‘dep. + hyp.’) and ‘no response’ for individual PNs. PNs are grouped based on their most frequent response type; depolarized (red, 36/54 PNs); hyperpolarized (blue, 6/54 PNs); depolarized + hyperpolarized (purple, 1/54 PN); no response (grey, 11/54 PNs), bars show mean and SD for all PNs in each group. (i) Averages of subthreshold voltage (red) and calcium (green) dendritic signaling during sensory stimulation (only recordings with no dendritic spikelets and both depolarizing and hyperpolarizing signals are included) and corresponding eye responses (blue), black lines show mean, colored traces show SD. (j) Relationship between average subthreshold dendritic voltage and calcium during the stimulus, solid line is linear regression (Pearson’s r = 0.71, p = 4.51×10^−10^) and dashed lines show 95% confidence intervals.

Throughout the experiment, fully awake mice were sitting on a rotating treadmill, head-fixed under a two-photon microscope (Figure 1d). Linescan recordings (10 seconds in length) included a 100ms sensory stimulus beginning 5 seconds after recording onset, giving an air puff directed at the ipsilateral eye. Air-puffs (pressure: 30psi) were delivered via a glass capillary positioned 2 cm from the eye to evoke a reliable eye blink reflex in the ipsilateral eye, which was monitored using a video camera (Figure 1e). This form of sensory stimulation was chosen to evoke dendritic calcium signals in PNs of lobule V, as described previously (Najafi et al., 2014a). Mice were naïve to the sensory stimulus receiving (< 5) test stimuli before recording began and 5-30 stimuli (average of 13 recordings per PN) during a single recording session. During sessions all air puffs were unexpected, triggering an eye blink reflex in the ipsilateral eye and no conditioned pre-stimulus eye blink responses were observed, indicating that the mouse did not learn to expect the stimulus (Figure 1 – figure supplement 1).

We previously described two types of suprathreshold dendritic signals in the PN dendrites of awake mice; dendritic complex spikes ‘DCS’ triggered by CF input and single dendritic spikes ‘DS’ triggered by strong PF input (Roome and Kuhn, 2018) (see Figure 1 – figure supplement 2 and Methods for DCS and DS detection criteria). DCS comprise a burst of rapid and distinct spikelets (typically 2-5 spikelets, each 1-2 ms in duration), associated with a large dendritic calcium signal. DS are characterized by a gradually ramping membrane potential, followed by a single spikelet and small associated calcium signal. We also described dendritic voltage signals evoked by synaptic input that correlate with action potential firing at the PN soma, but these did not trigger dendritic spikes and associated calcium signals, and so we refer to these as ‘subthreshold’ dendritic signals (Roome and Kuhn, 2018).

### Sensory stimulation evokes sub- and suprathreshold signals in Purkinje neuron dendrites of awake mice

Simultaneous voltage and calcium dendritic recordings revealed both subthreshold and suprathreshold signaling during sensory stimulation (Figure 1f). Suprathreshold signals (DCS and DS) were often, but not always, detected during coincident subthreshold signals. For example (Figure 1f), two PNs, PN1 and PN2, in the same mouse show responses; PN1 shows a slow subthreshold depolarization (black trace) with a DCS riding on top (red trace), while PN2 inset shows only a DCS, with no detectable subthreshold signal. Occasionally we also detected only subthreshold signals during sensory stimulation, with no coincident suprathreshold signal (see below). The sub- and suprathreshold dendritic signals were not evoked during voluntary eye blinks or detected while the mouse was anaesthetized and presented with the same air-puff stimuli (Figure 1 – figure supplement 3 and 4, respectively).

Sensory evoked subthreshold dendritic signaling has not been explored in awake animals. Thus, we began by analyzing the subthreshold dendritic signals. The amplitude and direction (i.e. depolarizing or hyperpolarizing) of evoked subthreshold voltage signals was variable between individual PNs and on a trial by trial basis (Figure 1 g-j and Figure 2a). By removing all suprathreshold signals (see Methods and Figure 1 – figure supplement 2 for spikelet detection and removal), we sorted voltage recordings based upon the amplitude and direction of the sensory evoked subthreshold response (Figure 1g and Figure 1 – figure supplement 2). Approximately 52% of voltage recordings displayed a depolarizing subthreshold signal (with detection threshold > 2 standard deviations above baseline), and 25% showed no response during the stimulus (n = 696). The remaining 14% showed hyperpolarizing voltage signals (detection threshold > 2 standard deviations below baseline) and 7% had a truncated signal that appeared to be a superposition of both depolarizing and hyperpolarizing signals (detection threshold > 2 standard deviations above and below baseline at different time points during the stimulus).

**Figure 2.**
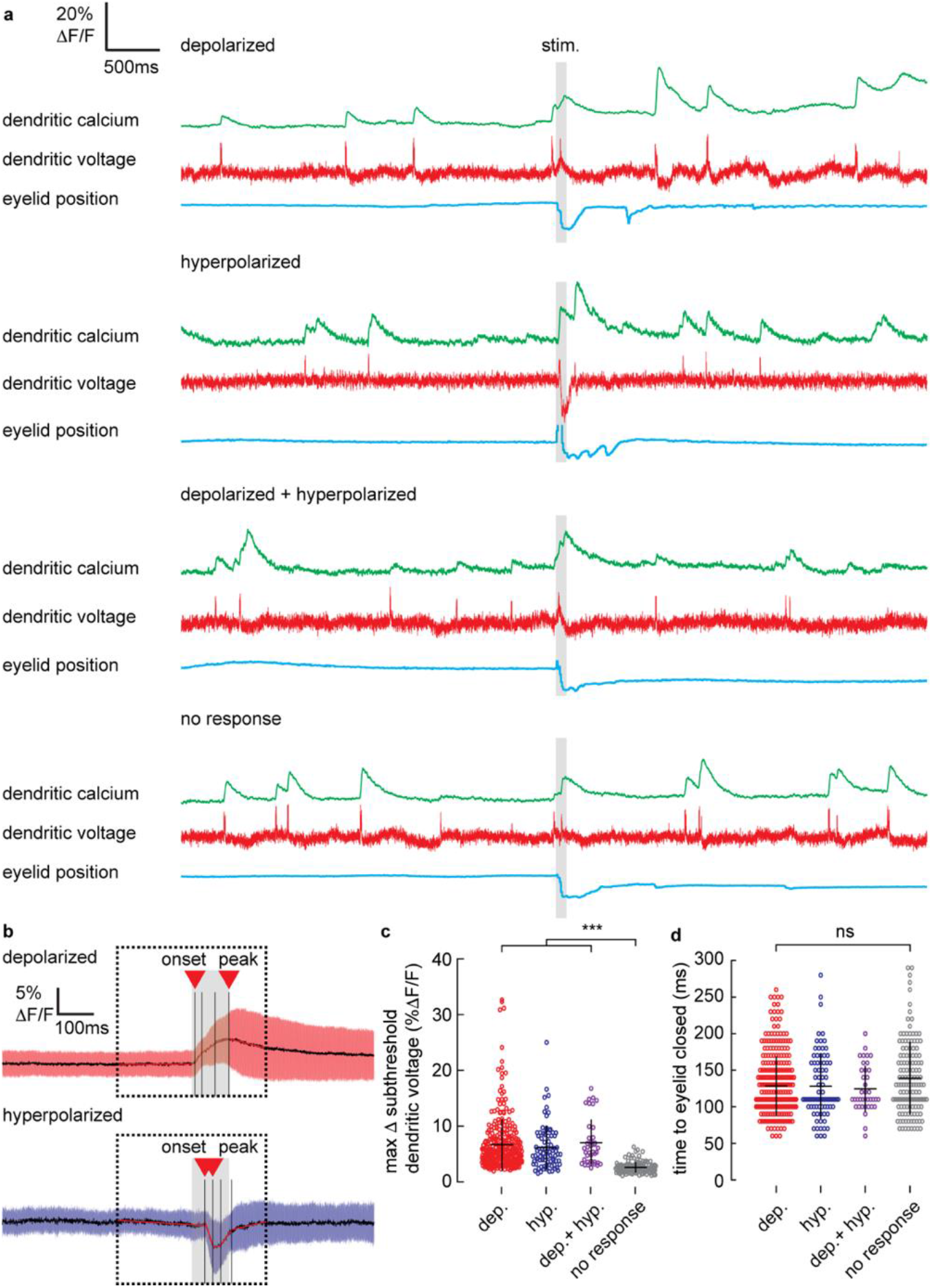
Stimulus evoked subthreshold dendritic voltage signals and their relationships with eyelid closure time. (a) Single trial recordings of simultaneous dendritic voltage (red traces) and calcium (green traces) during sensory stimulation (stim.). Suprathreshold signals are clearly visible in both voltage and calcium traces. Blue traces show average intensity of ROI used to record eyelid position (eyeblink responses). Grey bars show full duration of air puff stimulus. Each example shows subthreshold voltage signals during stimulus belonging to each response type; depolarized, hyperpolarized, depolarized + hyperpolarized or no response (top to bottom respectively). Recordings were sorted based on the subthreshold response during the stimulus, by first removing spikelets from the voltage recording, binning in 10ms epochs, and thresholding at ± 2 standard deviations. (b) Estimates for onset time and peak (shown by red triangles) in depolarizing (top) and hyperpolarizing (bottom) signals were calculated by minimizing the sum of residual (squared) error within four regions of the signal (shown by black bars) and fitting a linear regression within each region (red lines). (c) Maximum change in subthreshold voltage measured during the stimulus for all recordings in each response group. (d) Time for eyelid to close after stimulus onset for each response group. Horizontal and vertical bars show mean ± SD respectively. *** indicates p < 0.001.

Individual PNs displayed a trend towards a particular subthreshold voltage response (Figure 1h). The majority of PNs exhibited ‘depolarizing’ responses (36/54 PNs), and the remaining PNs exhibited either ‘hyperpolarizing’ (6/54 PNs), ‘no response’ (11/54 PNs), or both ‘depolarizing + hyperpolarizing’ (1/54 PNs). The average fraction of recordings across all PNs showing their preferred response was 0.65 ± 0.18 (mean ± SD, n = 54 PNs). Combining voltage recordings from all groups, we found that the average amplitude of all sensory evoked subthreshold voltage signals was positive relative to baseline (i.e. depolarizing); (2.91 ± 4.3% ΔF/F, mean ± SD, 54 PNs, t-test, p = 7.25×10^−6^), (Figure 1 – figure supplement 5). Therefore, on average, the sensory evoked subthreshold voltage signal is excitatory, and so elicited by PF synaptic input.

While dendritic calcium signals in PNs are predominantly suprathreshold and mediated by P/Q-type voltage gated calcium channels (VGCCs) (Usowicz et al., 1992), additional mechanisms also contribute to dendritic calcium signaling at PF-PN synapses, that do not require suprathreshold dendritic spikes. For example, glutamate released during PF input can activate group 1 metabotropic glutamate receptors (mGluR1), which are also expressed at PF-PN synapses, triggering calcium influx (Tempia et al., 2001) and modulating low threshold T-type VGCCs (Ait Ouares and Canepari, 2020; Hildebrand et al., 2009; Otsu et al., 2014). Whether or not these signals are involved in dendritic signaling during sensory stimulation has not been confirmed.

To determine if subthreshold dendritic calcium signals are evoked by sensory stimulation, we selected all recordings in which no suprathreshold (DCS or DS) signals were detected (i.e. no suprathreshold dendritic spikes and calcium signals) in a window ±300ms of the stimulus onset. Averaging these recordings revealed a gradual increase in dendritic calcium during sensory stimulation (dendritic calcium at stimulus offset relative to baseline: 1.8 ± 2.8 %ΔF/F, mean ± SD, n = 59, t-test, p = 5.9×10^−6^) (Figure 1i). Calculating the average subthreshold voltage and calcium signal over the full sensory stimulus, on a trial by trial basis, revealed a subthreshold voltage-calcium relationship (n = 59, Pearson’s r = 0.71, p = 4.51×10^−10^) (Figure 1j).

Single trial recordings of simultaneous dendritic voltage (red traces) and calcium (green traces), during sensory stimulation (stim.) for each subthreshold voltage response type; ‘depolarizing’, ‘hyperpolarizing’, ‘depolarizing + hyperpolarizing’ and ‘no response’, are shown in Figure 2a. The temporal profile of each voltage response was resolved by sorting and averaging voltage signals in each response group (Figure 2b). Depolarizing signals increased gradually throughout the stimulus, beginning 9 ± 3 ms (mean ± SD, n = 307) after stimulus onset, and reaching a maximum after 101 ± 3 ms. Hyperpolarizing signals were sharper, and their onset occurred later, beginning 35 ± 2 ms (mean ± SD, n = 83) after stimulus onset, and reaching a minimum 57 ± 2 ms (mean ± SD) after stimulus onset.

We calculated the maximum change in subthreshold voltage during the sensory stimulus (excluding dendritic spikes), for all recordings in each response group (Figure 2c). The maximum change in subthreshold voltage was similar between depolarizing (6.7 ± 4.7 %ΔF/F, mean ± SD, n = 307), hyperpolarizing (6.1 ± 4.0 %ΔF/F, mean ± SD, n = 83) and depolarizing + hyperpolarizing groups (7.0 ± 4.2 %ΔF/F, mean ± SD, n = 38) and significantly larger than the no response group (2.6 ± 1.0 %ΔF/F, mean ± SD, n = 133), ANOVA, p < 1.2×10^−8^, n = 561, 49 PNs. The time of eyelid closure (measured from stimulus onset to maximum eyelid closure) was independent of response type (Figure 2d); depolarized (128 ± 40 ms, mean ± SD), hyperpolarized (123 ± 44 ms, mean ± SD), depolarized + hyperpolarized (124 ± 32 ms, mean ± SD) and no response (139 ± 50 ms, mean ± SD) groups, Kruskal-Wallis ANOVA, p > 0.35, n = 561, 49 PNs. This confirmed that a lack of subthreshold response was not due to a failure of the sensory stimulus.

We used pharmacology to investigate the origin of the stimulus evoked dendritic signals. All sub- and suprathreshold signals were blocked by AMPA receptor antagonist CNQX (100μM) (and also by Na^+^ channel blocker Lidocaine (0.2%), Figure 3 – figure supplement 1). Single trial recordings of simultaneous dendritic voltage and calcium signaling during sensory stimulation under control and CNQX conditions from the same PN are shown in Figure 3a. Average eyelid responses under control (n = 45) and CNQX (n = 37) conditions from 5 PNs are shown in Figure 3b. The eyelid closure time following CNQX application was not significantly different to control conditions (Figure 3c); control (134 ± 40 ms, mean ± SD); CNQX (156 ± 44 ms, mean ± SD), Kolmogorov-Smirnov, p = 0.12, n = 82, 5 PNs.

**Figure 3.**
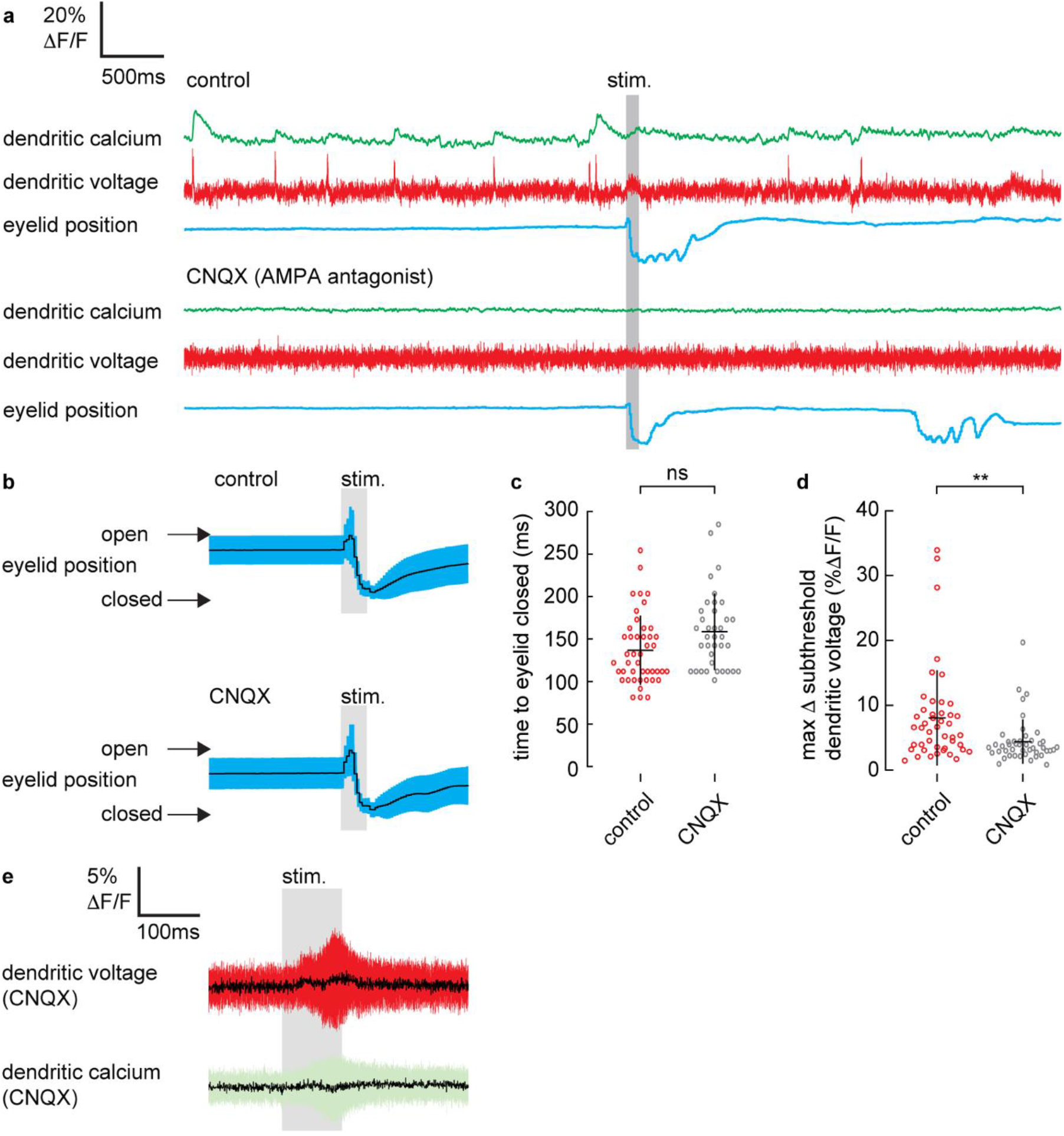
AMPA receptor antagonist, CNQX, blocks stimulus evoked subthreshold dendritic voltage and calcium signaling. (a) Single trial recordings of simultaneous dendritic voltage (red traces) and calcium (green traces) during sensory stimulation (stim.). Blue traces show average intensity of ROI used to record eyelid position (eyeblink responses). Grey bars show full duration of air puff stimulus. Top example shows voltage and calcium recordings during control conditions. Note in this example sensory stimulation evoked only a subthreshold voltage signal and no suprathreshold (calcium) signals during the stimulus. Bottom example shows recordings following CNQX application from the same PN. (b) Mean eye responses from control (top; n = 45) and CNQX (bottom; n = 37) recordings (black trace shows mean; blue trace shows SD). (c) Time for eyelid to close after stimulus onset under control (red) and CNQX (grey) conditions. (d) Maximum change in subthreshold voltage measured within a 200ms window after stimulus onset under control and CNQX conditions. (e) Averages of subthreshold voltage (red) and calcium (green) dendritic signaling during sensory stimulation, following CNQX application, black line shows mean, colored trace shows SD. ** p < 0.01.

The maximum change in subthreshold voltage (measured within a 200ms window after stimulus onset), was significantly reduced under CNQX conditions; control (8.0 ± 7.3 % ΔF/F, mean ± SD, n = 45), and CNQX (4.4 ± 3.4 % ΔF/F, mean ± SD, n = 46), unpaired t-test, p = 0.003, n = 91, 5 PNs (Figure 3d). Accordingly, the voltage dependent subthreshold calcium signal we detect during sensory stimulus (described in Figure 1i-j), was also blocked by CNQX (Figure 3e) and also by Lidocaine (Figure 3 – figure supplement 1g); average calcium at stimulation offset under CNQX conditions was (−0.002 ± 0.023 % ΔF/F, mean ± SD) and not significantly different from baseline, t-test, p = 0.59, n = 36, 5 PNs.

Metabotropic γ-aminobutyric acid type B (GABA_B_) receptors are expressed at PF-PN synapses (Kulik et al., 2002), and so likely to contribute to inhibitory synaptic input at our recording location of the PN spiny dendrites. We assessed their contribution by applying GABA_B_ receptor antagonist, CGP 35348 (100μM). Single trial recordings of simultaneous dendritic voltage and calcium signaling during sensory stimulation under control and CGP conditions from the same PN are shown in Figure 4a. Average eyelid responses under control (n = 50) and CGP (n = 59) conditions from 5 PNs are shown in Figure 4b. Application of CGP reduced sensory evoked hyperpolarizing voltage signals, but not the depolarizing voltage signals. Average subthreshold voltage (with spikelets removed) under control (n = 50) and CGP (n = 59) conditions are shown in Figure 4c. Note that GABA_B_ blockade revealed a sustained sensory evoked depolarizing voltage signal, that continued after the stimulus had ended. The eyelid closure time following CGP application was not significantly different to control conditions (Figure 4d); control (114 ± 23 ms, mean ± SD, n = 50); CGP (125 ± 34 ms, mean ± SD, n = 59), Kolmogorov-Smirnov, p = 0.29, n = 109, 5 PNs. The maximum or minimum subthreshold dendritic voltage within a 200ms window after stimulus onset, under control and CGP conditions is shown in Figure 4e. Stimulus evoked subthreshold voltage signals were significantly more positive (depolarizing) following CGP application; control (−0.4 ± 6.0 % ΔF/F, mean ± SD, n = 50), and CGP (3.5 ± 7.7 % ΔF/F, mean ± SD, n = 59), Kolmogorov-Smirnov, p = 0.005, n = 109, 5 PNs.

**Figure 4.**
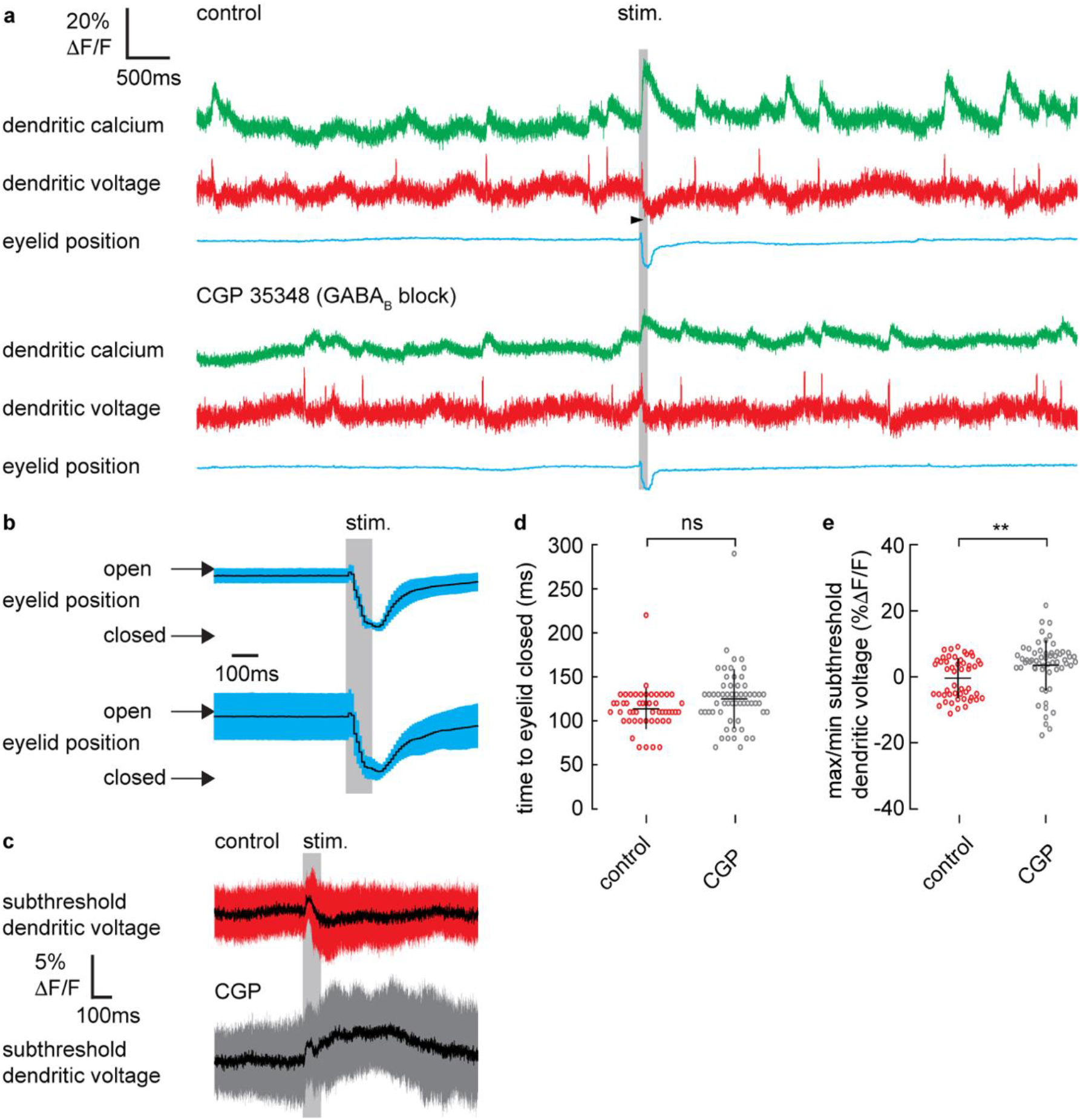
GABA_B_ receptor antagonist, CGP 35348, blocks stimulus evoked hyperpolarizing dendritic voltage signaling. (a) Single trial recordings of simultaneous dendritic voltage (red traces) and calcium (green traces) during sensory stimulation (stim.). Blue traces show average intensity of ROI used to record eyelid position (eyeblink responses). Grey bars show full duration of air puff stimulus. Top example shows voltage and calcium recordings during control conditions (black arrow indicates a hyperpolarizing signal towards the end of the stimulus), and bottom example shows recordings following CGP application from the same PN. Note there was a relative increase in baseline GCaMP6f fluorescence of 93 ± 90 % ΔF/F, following CGP application. (b) Mean eye responses from control (top) and CGP (bottom) recordings (black trace shows mean; blue trace shows SD). (c) Average subthreshold voltage (spikelets removed) under control (n = 50) and CGP (n = 59) conditions. Note that GABA_B_ blockade reveals a sustained sensory evoked depolarizing voltage (black traces show mean; colored traces show SD). (d) Time for eyelid to close after stimulus onset under control (red) and CGP (grey) conditions. (e) Maximum or minimum subthreshold voltage measured within a 200ms window after stimulus onset under control and CGP conditions. ** p < 0.01.

Sensory evoked stimulation has previously been shown to trigger enhanced calcium signals in PN dendrites of the cerebellar vermis, compared to spontaneous calcium signals (Najafi et al., 2014b). Various mechanisms have been proposed to modulate suprathreshold dendritic calcium signals in PNs, including molecular layer interneuron activity (Callaway et al., 1995; Gaffield et al., 2018; Kitamura and Häusser, 2011), granule cell activity (Najafi et al., 2014a, b) and graded CF input (Gaffield et al., 2019). However, the dendritic suprathreshold voltage signals that trigger these calcium signals have not been directly explored in awake animals. Thus, we next examined suprathreshold dendritic voltage signaling evoked by sensory stimulation (Figure 5).

**Figure 5.**
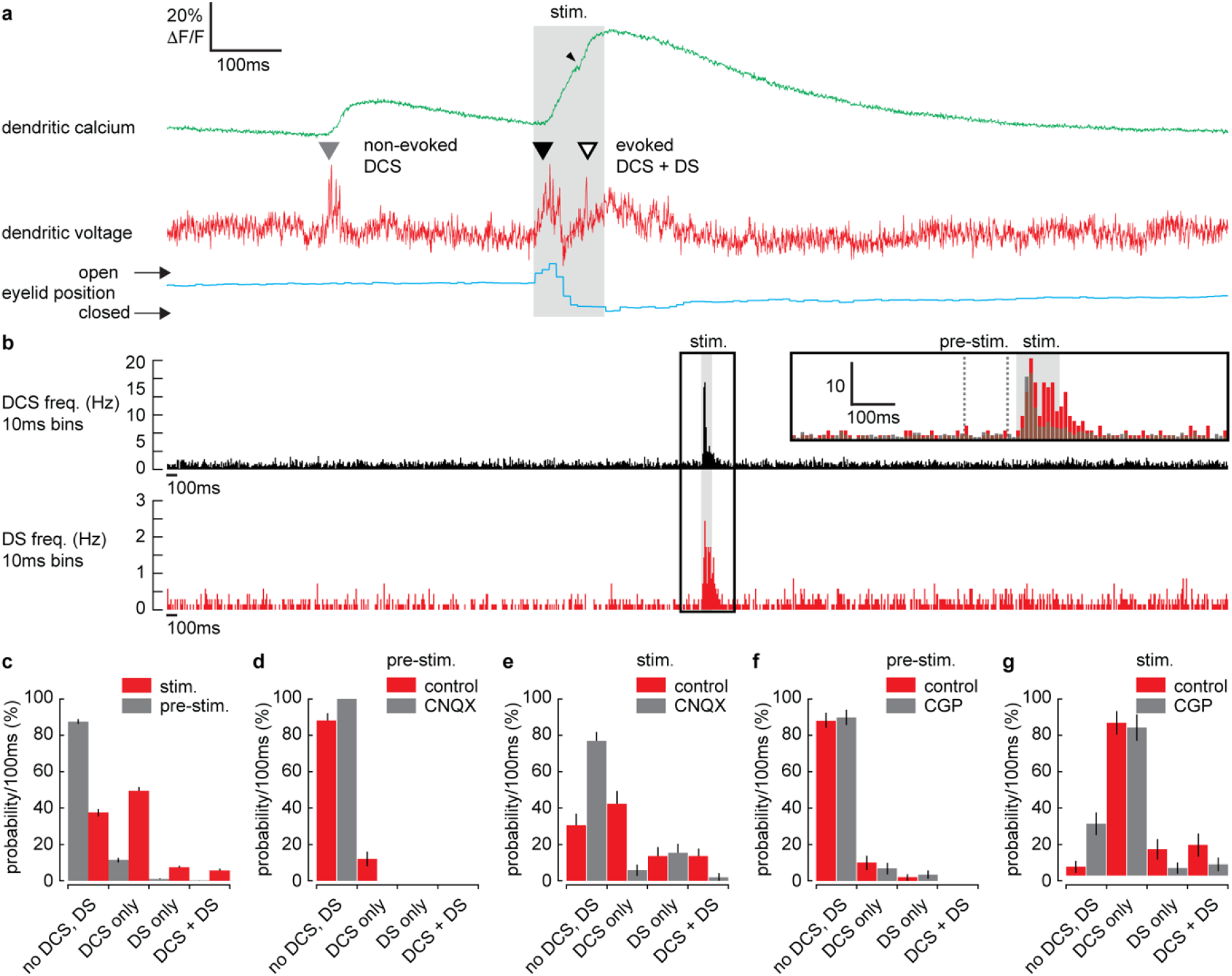
Suprathreshold calcium spikes are triggered by CF and PF synaptic input during sensory evoked responses. (a) A simultaneous recording of dendritic calcium (green trace) and voltage (red trace) during air puff sensory stimulation, showing DCS (filled triangle) and DS (open triangle) signals. Signals that occur during the air puff stimulus (stim.) are termed ‘evoked’ (black triangle), otherwise ‘non-evoked’ (grey triangle). Small black arrowhead indicates increase in calcium signal following the DS. Corresponding eyelid response (blue trace), timing of air puff stimulus (gray bar). (b) PSTH for DCS frequency (top black) and DS frequency (bottom red) using 10ms bins. Inset shows zoomed overlay of DCS (black) and DS (red) frequencies, relative to their average frequencies. Dashed grey lines indicate the pre-stimulus period (100ms window prior to air puff onset). (c) Probability distributions for all possible outcomes; ‘no DCS, DS’, ‘DCS only’, ‘DS only’, and paired ‘DCS + DS’, during pre-stimulus and stimulus time windows. Probability distributions for suprathreshold DCS and DS signals during pre-stimulus (d) and stimulus (e) time windows, under control and CNQX conditions. Probability distributions for suprathreshold DCS and DS signals during pre-stimulus (f) and stimulus (g) time windows, under control and CGP conditions. Vertical bars in probability distributions show bootstrapped SD.

As described previously (Roome and Kuhn, 2018), we detected two types of suprathreshold signals in the PN spiny dendrites; CF evoked DCS, which occurred at a reliable average rate of 1.05 ± 0.4 Hz (mean ± SD, 54 PNs, 696 recordings) and PF evoked DS, which occurred irregularly at an average rate of 0.13 ± 0.3 Hz (mean ± SD, 54 PNs, 696 recordings. Importantly, we detected an increase in frequency of both DCS and DS suprathreshold signals during the sensory stimulus (Figure 5a). Peri-stimulus time histograms (PSTH) for suprathreshold DCS and DS signals (Figure 5b), show how their frequency increased dramatically during the sensory stimulus. Average DCS and DS frequencies during the stimulus was 5.73 ± 5.51 Hz and 1.24 ± 0.77 Hz (mean ± SD), respectively; a relative increase in DCS and DS frequency of 5.5 and 9.5 respectively (see Figure 5b inset). The average timing of DCS and DS signals after stimulus onset was 43.3 ± 22.1 ms (mean ± SD, n = 414) and 53.3 ± 24.5 ms (mean ± SD, n = 92) respectively.

We compared suprathreshold signals in a pre-stimulus time window (100ms window prior to stimulus onset, see Figure 5b inset) with the stimulus time window, in all recordings and calculated the probability for the following outcomes; ‘no DCS, DS’, ‘DCS only’, ‘DS only’, and paired ‘DCS + DS’ (Figure 5c). There was a dramatic increase in the probability of detecting suprathreshold signals during the stimulus compared to the pre-stimulus time window; DCS only (pre-stimulus: 11.5 ± 1.2 %, stimulus: 49.3 ±1.7 %); DS only (pre-stimulus: 0.8 ± 0.4 %, stimulus: 7.6 ± 1.1 %); paired DCS + DS signals (pre-stimulus: 0.3 ± 0.2 %, stimulus: 5.9 ± 0.9 %) (mean ± SD, n = 688, Kolmogorov-Smirnov, p = 7.3×10^−79^). CNQX application blocked suprathreshold DCS and DS signals. Probability distributions for suprathreshold DCS and DS signals during pre-stimulus time windows and during the stimulus, under control and CNQX conditions are shown in Figure 5d and Figure 5e, respectively. There were no suprathreshold signals detected during the pre-stimulus time window following CNQX application, and probability distributions were shifted towards a higher probability of ‘no-signal’ following CNQX application during the stimulus time window, Kolmogorov-Smirnov, p = 1.6×10^−5^, n = 91, 5 PNs. Conversely, CGP application did not significantly alter the probability of suprathreshold signals, during either pre-stimulus or stimulus time windows (Figure 5f and Figure 5g, respectively), Kolmogorov-Smirnov, pre-stimulus; p = 1.0, stimulus; p = 0.22, n = 109, 5 PNs.

In agreement with previous studies (Gaffield et al., 2019; Najafi et al., 2014b), we found that average evoked calcium signals (i.e. signals triggered during the stimulus) were enhanced relative to non-evoked calcium signals (i.e. signals occurring before the stimulus), (relative calcium peak; 1.4 ± 0.6, 51 PNs, 365 recordings, t-test, p = 1.5×10^−5^), (Figure 1 – figure supplement 5). To determine the mechanisms underlying the enhanced calcium signals, it is necessary to consider the suprathreshold voltage signals that trigger them.

To do this we examined the individual suprathreshold signals. Suprathreshold dendritic voltage spikelets and calcium signals detected during the stimulus (termed ‘evoked’ signals), were compared with signals detected prior to the stimulus (termed ‘non-evoked’ signals). We counted the total number of spikelets detected in a 100ms window following each evoked or non-evoked DCS signal, including any additional DCS and DS signals within that time window, and measured the resulting calcium signal peak. Calcium peaks were on average larger following evoked DCS signals compared to the non-evoked DCS signals (non-evoked: 12.6 ± 9.3 %ΔF/F, n = 1624, evoked: 20.8 ± 15.1 %ΔF/F, n = 333, mean ± SD, unpaired t-test, p = 1.6×10^−37^) (Figure 6a). Accordingly, the corresponding dendritic voltage recordings showed an increase in the total number of dendritic spikelets following each evoked DCS signal compared to non-evoked DCS signals, evident through a positive shift in spikelet count probability distribution (Kolmogorov-Smirnov, p = 1.85×10^−5^) (Figure 6b).

**Figure 6.**
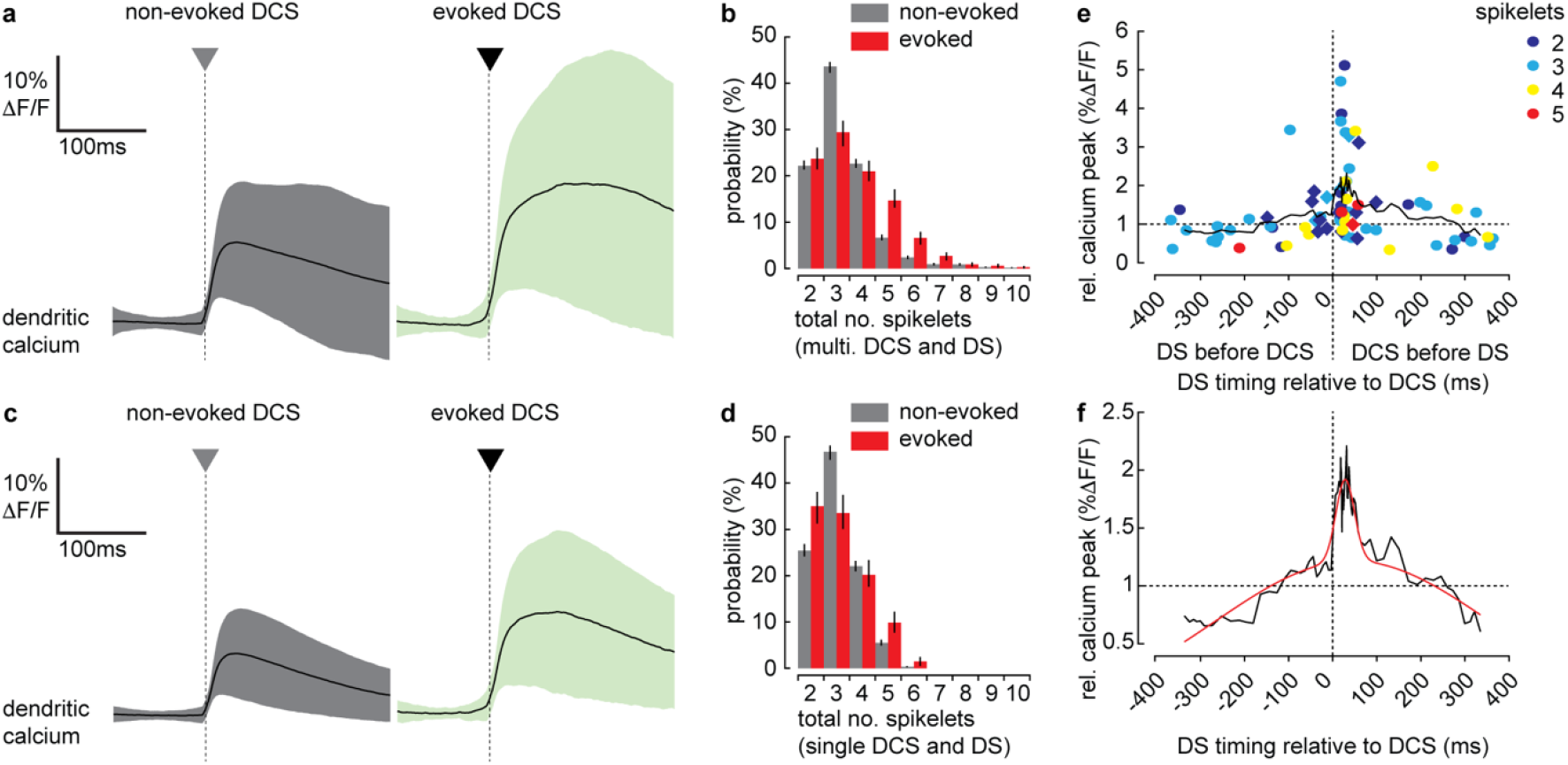
Supra-linear relationship between the timing of DCS and DS signals and their corresponding calcium signals. (a) Averages of all calcium signals following non-evoked (grey, left) and evoked (green, right) DCS signals. Black trace is mean and colored trace is SD. (b) Probability distributions for the total number of dendritic spikelets detected in a 100ms window following onset of each non-evoked and evoked DCS signal. (c) Averages of all calcium signals following non-evoked (grey, left) and evoked (green, right) DCS signals, excluding multiple DCS signals. Black trace is mean and colored trace is SD. (d) Probability distributions for the total number of dendritic spikelets detected in a 100ms window following onset of each non-evoked and evoked DCS signal, excluding multiple DCS signals. (e) Relationship between DCS + DS pair timing and the corresponding relative calcium peak. Relative calcium peaks were calculated with respect to the average calcium signal of DCS with the same number of spikelets (but with no paired DS signal). Colors represent the number of DCS spikelets for each DCS + DS pair. Data points for evoked DCS + DS pairs are diamonds and non-evoked DCS + DS pairs are circles. Black line shows boxcar average of 10 consecutive data points and (f) two-term gaussian fit (red line) applied to boxcar average (adjusted r-square = 0.9117). Vertical bars in probability distributions show bootstrapped SD.

We considered the possibility that the increase in dendritic calcium peak and the number of spikelets result from an increase in the number of CF inputs (i.e. multiple DCS signals) within the 100ms time window. Due to slow calcium indicator dynamics, high frequency CF inputs cannot always be resolved from the calcium signal alone (see Figure 1 – figure supplement 2c). Using the voltage recording we detected and excluded all signals in which multiple CF inputs were observed and repeated the analysis. Interestingly, when only single DCS signals were permitted, the peak calcium of evoked signals remained enhanced compared to non-evoked signals (non-evoked: 11.9 ± 8.1 %ΔF/F, n = 1292, evoked: 19.7 ± 14.2 %ΔF/F, n = 203, mean ± SD, unpaired t-test, p = 3.8×10^−28^) (Figure 6c). However, the difference between pre-stimulus and stimulus probability distributions of spikelet numbers was no longer significant (Kolmogorov-Smirnov, p = 0.075) (Figure 6d), indicating that the enhanced calcium signal was not exclusively due to an increase in CF input frequency, or even due to an increase in dendritic spiking, caused by stronger CF input alone.

Since both DCS and DS frequency increased during the sensory stimulus, we next considered the possibility that enhanced evoked calcium signals result from paired DCS and DS signals that occur during the same stimulus. We investigated the relationship between the timing of paired DCS + DS signals and their corresponding dendritic calcium signal, by selecting all DCS + DS pairs (non-evoked and evoked) with < 400ms temporal separation. We measured the calcium peak following each pair of DCS + DS signals, relative to the average calcium peak of a single DCS signal comprising the same number of spikelets. Relative calcium peaks were plotted in relation to the temporal separation of each DCS + DS pair (Figure 6e). Analysis revealed a clear asymmetric and supra-linear relationship between the timing of DCS and DS signals and their corresponding calcium signals. By fitting a two-term gaussian function (Figure 6f) we estimate that the average relative calcium peak is maximum when DS follow DCS signals by 28.3 ± 5.4 ms (mean ± SD).

Thus, we confirm that paired DCS + DS signals evoked by coincident CF and PF input act to enhance the dendritic calcium signal. However, paired DCS + DS signals are relatively rare (detected in only 5.9 ± 0.9 % of recordings), suggesting an additional coincidence detection mechanism at work. Taken together, our findings thus far reveal an increase in both sub- and suprathreshold dendritic signaling during sensory stimulation. These signals originate primarily from coincident PF and CF synaptic input, because the subthreshold signals are predominantly depolarizing, and suprathreshold dendritic signals are triggered exclusively by excitatory CF and PF synaptic input.

### Supra-linear dendritic signaling in PN dendrites during sensory evoked stimulation is voltage and mGluR1 dependent

To investigate the mechanisms underlying evoked supra-linear calcium signals further, we selected individual evoked and non-evoked DCS signals only (now with no subsequent DCS or DS signals). We sorted DCS signals based on the number of spikelets generated. This created four DCS groups, ranging from 2 to 5 spikelets) (Figure 7a-c). Averaging voltage traces within each evoked spikelet group revealed a prominent subthreshold depolarization beginning before the evoked DCS signals, this ‘pre-DCS’ signal was not detected preceding non-evoked DCS signals (black arrowheads in Figure 7c). This would be expected because on average, sensory evoked subthreshold signals are depolarizing and precede DCS generation (as shown in Figure 2 and Figure 6). Average pre-DCS voltages (measured in a 10 ms window preceding DCS signals) for each group are shown in Figure 7d (evoked pre-DCS vs. non-evoked pre DCS, ANOVA, p < 0.004). The evoked pre-DCS depolarization amplitude was independent of number of DCS spikelets (2 spikelets: 1.31 ± 1.84 % ΔF/F; 3 spikelets: 1.35 ± 1.93 % ΔF/F; 4 spikelets: 1.47 ± 2.22 % ΔF/F; 5 spikelets: 1.29 ± 1.14 % ΔF/F, mean ± SD, ANOVA, p > 0.52) (Figure 7e).

**Figure 7.**
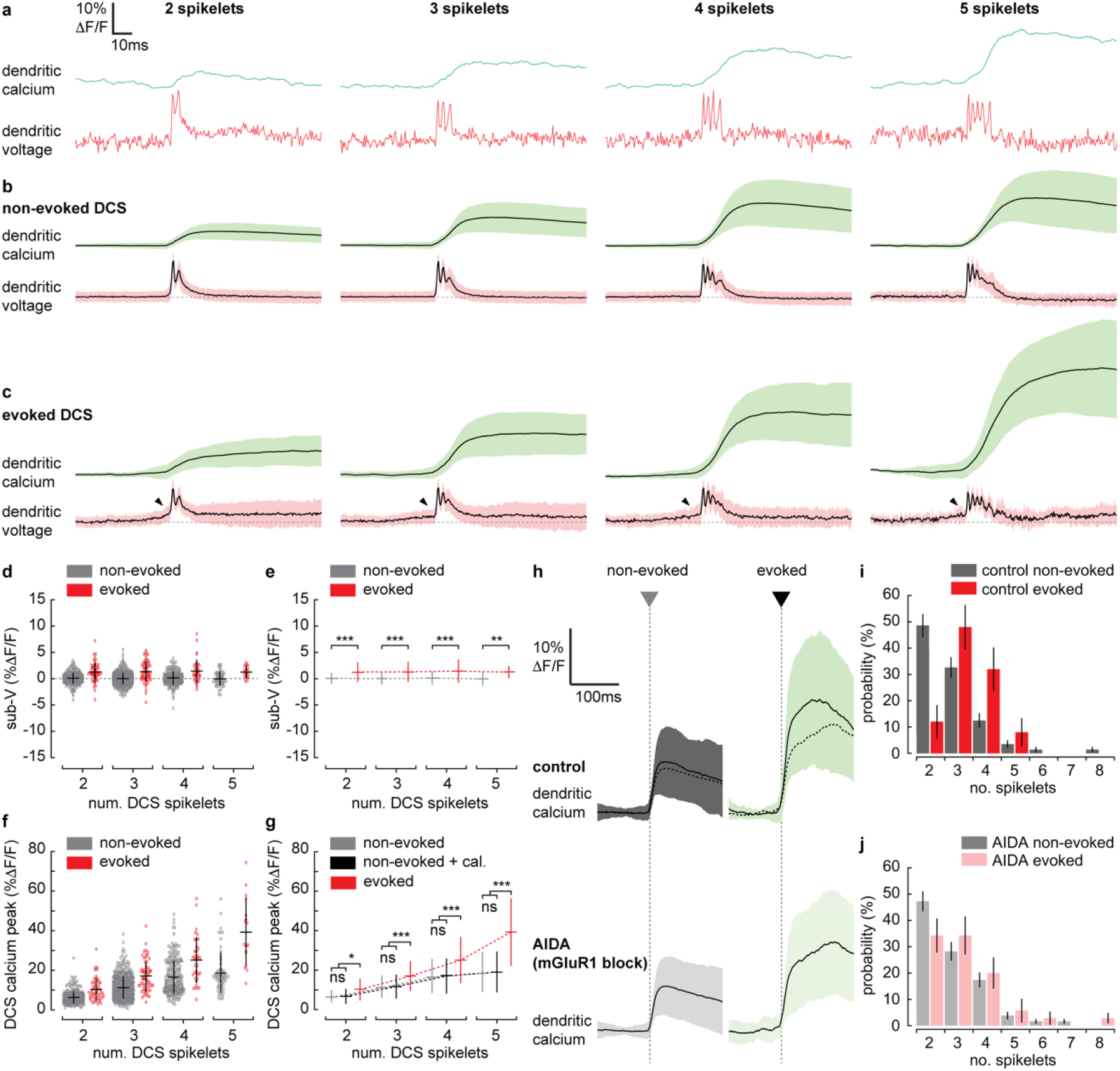
Supra-linear dendritic signaling in PN dendrites during sensory evoked stimulation is voltage and mGluR1 dependent. (a) Single trial examples of non-evoked DCS signals sorted based on the number of spikelets showing dendritic voltage (red traces) and calcium (green traces). (b) Averages of all non-evoked DCS signals. (c) Averages of all evoked DCS signals. Arrowheads indicate subthreshold depolarization preceding evoked DCS signals. (d) Average subthreshold pre-DCS voltage for evoked (red) and non-evoked (grey) signals, grouped based on the number of DCS spikelets. Horizontal bars show mean, vertical bars show SD. (e) Mean from all groups in (d). (f) Average calcium peak for evoked (red) and non-evoked (grey) signals, grouped based on the number of DCS spikelets. (g) Mean from all groups in (f) and sum of non-evoked and + cal. groups (black). (h) Averages of all calcium signals following non-evoked (grey, left) and evoked (green, right) DCS signals, under control (top dark traces) and mGluR1 block (AIDA) conditions (bottom light traces and top overlaid dashed traces). Black trace is mean and colored trace is SD. Probability distributions for the total number of dendritic spikelets detected in a 100ms window following onset of each non-evoked and evoked DCS signal, excluding multiple DCS signals, under (i) control and (j) mGluR1 block conditions (Kolmogorov-Smirnov, vertical bars in probability distributions show bootstrapped SD). * p < 0.05, ** p < 0.01 and *** p < 0.001.

We would expect the subthreshold depolarization detected during evoked signals to contribute to the calcium signal through the voltage-dependent increase in calcium baseline that we described in Figure 1j. Using the sub-threshold voltage-calcium relationship determined in Figure1j, and the average pre-DCS voltage across all spikelet groups; 1.36 ± 1.89 % ΔF/F (approximately 2.7 ± 3.8 mV, mean ± SD), we would predict an average increase in calcium baseline of 0.8% ± 1.5 % ΔF/F.

In reality, the average calcium peak was much larger during evoked DCS signals compared to non-evoked DCS signals comprising the same number of spikelets. Furthermore, the difference between evoked and non-evoked calcium peaks increased supra-linearly with the number of DCS spikelets (non-evoked vs. evoked, 2 spikelets: 6.2 ± 3.2 % ΔF/F vs. 10.1 ± 5.5 % ΔF/F; 3 spikelets: 11.1 ± 5.7 % ΔF/F vs. 17.0 ± 7.7 % ΔF/F; 4 spikelets: 16.4 ± 8.6 % ΔF/F vs. 25.1 ± 11.7 % ΔF/F; 5 spikelets: 18.5 ± 10.2 % ΔF/F vs. 39.3 ± 17.1 % ΔF/F, mean ± SD, ANOVA, p < 0.002) (Figure 7f). In this case, the linear sum of non-evoked DCS calcium peak and our predicted increase in calcium baseline (‘+ cal.’) could not account for the supra-linear enhancement in calcium peak during evoked DCS signals (non-evoked + cal. vs. evoked, ANOVA, p < 0.043) (Figure 7g).

Activation of mGluR1s by coincident PF and CF input is known to drive supra-linear calcium signals at PF-PN synapses in cerebellar slices (Wang et al., 2000), but their contribution to PN dendritic signaling has not been explored in vivo. By application of AIDA (a potent and selective mGluR1 antagonist), we assessed the contribution of mGluR1 to sensory evoked dendritic signals in the awake mouse. After application of AIDA, sensory evoked calcium signals were significantly reduced (control: 24.4 ± 12.3 % ΔF/F, AIDA: 17.9 ± 9.3 % ΔF/F, mean ± SD, unpaired t-test, p = 0.025) (Figure 7h). Non-evoked calcium signals however were not significantly altered (control: 10.6 ± 6.5 % ΔF/F, AIDA: 9.5 ± 5.5 % ΔF/F, mean ± SD, unpaired t-test, p = 0.116), indicating that the contribution of mGluR1 activity to the calcium signal is specific to sensory evoked calcium signals when CF and PF inputs coincide, and we propose that mGluR1s are activated at PF-PN synapses, where the PF and CF synaptic potentials converge.

Probability distributions for suprathreshold signals detected in pre-stimulus and stimulus time windows were calculated under control and AIDA conditions. As in Figure 5c, there was a clear increase in signal probability during the stimulus compared to pre-stimulus time windows under both control and AIDA conditions, but we detected no difference between control vs. AIDA conditions (for either pre-stimulus or stimulus groups), indicating that mGluR1 block did not directly affect CF and PF input (Figure 7 – figure supplement 1). There was also no significant effect of AIDA on evoked subthreshold signaling, supporting the conclusion that the subthreshold signals are predominantly AMPA mediated (Figure 7 – figure supplement 1).

However, we found that the probability distribution for the total number of spikelets following evoked DCS was shifted towards lower numbers under AIDA conditions. Consequently, unlike control conditions (Figure 7i), the probability distributions for the total number of spikelets following evoked and non-evoked DCS signals were no longer significantly different following AIDA application (Figure 7j). This suggests that mGluR1 activity enhances spikelet generation during evoked DCS signals. Such an interaction between mGluR1 activity and spikelet generation, has previously been postulated (Tempia et al., 2001) and is likely mediated through their interaction with T and P/Q-type VGCCs (Hildebrand et al., 2009; Otsu et al., 2014).

## Discussion

Dendritic coincidence detection is a salient signal in neuronal processing. We used simultaneous voltage and calcium two-photon imaging of single PN dendrites at high temporal resolution (2kHz) to explore PF and CF coincident input during sensory evoked responses. The ability to distinguish between sub- and suprathreshold signaling and dendritic spikes evoked by either CF or PF input has provided fresh insight into graded dendritic calcium signals evoked by sensory stimulation, and the integration of information transferred by coincident PF and CF input pathways.

Here we describe how PF and CF synaptic input to PN dendrites is integrated during sensory evoked stimulation in awake mice. To summarize, we identify three interplaying dendritic coincidence detection mechanisms that modulate calcium signals at the PN dendritic level. PN dendrites integrate PF and MLI evoked excitatory and inhibitory post-synaptic potentials, resulting in a subthreshold voltage-dependent calcium signal (Figure 1). The first mechanism of dendritic coincidence detection occurs if the subthreshold voltage-dependent calcium signal (Figure 1j) coincides with a suprathreshold input. This is potentially the cause of the enhanced evoked calcium signals compared to non-evoked signals, while mGluR1 is blocked (Figure 7h). Previous in vitro studies have shown how membrane potential and timing of coincident sub- and suprathreshold synaptic input is critical for modulating calcium signaling and synaptic plasticity at PF-PN synapses (Ly et al., 2016; Sarkisov and Wang, 2008). The second mechanism is through PF evoked DS, whose frequency increases during the sensory stimulus and when paired with DCS, act to enhance the CF evoked calcium signal in a time-dependent manner (Figure 6). It is important to note that while PF evoked DS signals are relatively rare in awake mice (Roome and Kuhn, 2018), we found that their frequency increased 9.5 times during sensory stimulation, and their impact on the dendritic calcium signal was supra-linear. Thus, we propose PF evoked DS signals play a unique role in modulating dendritic calcium signals during sensory stimulation. In fact, the additional PF evoked DS could be viewed as a correction to the CF evoked DCS signal, supplementing an otherwise ‘too weak’ DCS signal with an extra dendritic spike. The third is through mGluR1 activity (Figure 7), which is known to trigger supra-linear calcium signals when PF and CF inputs are conjunctively activated, that modify PF-PN synaptic strength (Sarkisov and Wang, 2008; Wang et al., 2000). It is intriguing that the coincidence detection processes and supra-linear dendritic calcium signals we describe here, occur seemingly continuously; during sensory stimulation in awake naïve mice, and not only during motor learning of a specific task, such as eyeblink conditioning.

PN dendrites and their spines are highly sensitive to calcium levels, which can in turn determine the direction of synaptic plasticity at PF-PN synapses (Coesmans et al., 2004; Jorntell and Hansel, 2006). We expect that to constantly finetune plasticity at PF-PN synapses, calcium signals are continuously graded at the PN dendrite level, to permit a full dynamic range. The coincident activity of CF and PF synaptic input (and also PF driven feed forward inhibition via MLIs) combined with intrinsic dendritic mechanisms (VGCCs and mGluR1s) allow for this high dynamic range in calcium signaling. Here, we establish that their combined activity becomes more apparent during heightened sensory input, as indicated by their enhanced cooperation during sensory evoked eyeblink responses.

## Methods

### Animals and surgery

All animal procedures were conducted in accordance with guidelines of the Okinawa Institute of Science and Technology Institutional Animal Care and Use Committee in an Association for Assessment and Accreditation of Laboratory Animal Care (AAALAC International)-accredited facility. Cerebellar chronic cranial window surgeries were performed on 2-month-old male C57/BL6 mice, using a 5 mm glass cover slip with silicone access port (Roome and Kuhn, 2014). The window was positioned to allow imaging within lobule V of the cerebellar vermis and the access port was positioned to allow access to the imaging area via a micropipette for PN labelling and drug delivery.

### Microscope setup

We used a custom-built combined wide-field, two-photon microscope (MOM, Sutter Instruments) with either a ×5/N.A. 0.13 air objective (Zeiss) or a ×25/N.A. 1.05 water immersion objective with 2 mm working distance (Olympus) with ScanImage software. Bright field imaging was performed using a sCMOS camera (PCO.edge, PCO). A femtosecond-pulsed Ti:sapphire laser (Vision II, Coherent), circularly polarized and under-filling the back focal plane of the ×25 objective, was used to excite fluorescence (typical laser power at 1020 nm: 60 mW), which was detected by two GaAsP photomultiplier tubes (Hamamatsu) in the spectral range of 490–550nm (green: GCaMP6f) and 550–750 nm (red: ANNINE-6plus).

The mice were headfixed on a platform that consisted of a vertically rotating treadmill, head-plate stage and micromanipulator tower (Sutter Instruments), all mounted on a horizontally rotating stage (8MR190-90-4247, Standa). The sCMOS camera and infrared light source were used to record behavioral activity during recording at 100 fps. A second infrared video camera (Sony) was used to monitor mouse behavior throughout the experiment. The micromanipulator (M-285, Sutter instruments) was used for GCaMP6f virus (UPenn Vector Core) and ANNINE-6plus dye (www.sensitivefarbstoffe.de, Dr. Hinner and Dr. Hübener Sensitive Dyes GbR) injection and also for drug delivery and electrophysiological recording via micropipette.

### AAV injections

One week following surgery, mice were anesthetized (1-2% Isoflurane) and head mounted for two-photon guided injection of the adeno-associated viral vectors (AAVs) into the PN layer approximately 150 μm below the dura. For this, beveled quartz electrodes (0.7 mm ID, pulled and beveled to 10-20 μm tip diameter) containing AAV1.hSyn.*Cre* (2E13 GC/ml), AAV1.CAG.Flex.*GCaMP6f* (1.3E13 GC/ml), and 50 μM FITC in PBS at a ratio of 1:1:1 were used to specifically target PNs and visually control the position and size of the viral injection, while <0.1 PSI pressure was used to inject the virus for 1 min. After virus injection the pipette was retracted, and the mouse was returned to its cage.

### Single neuron labelling with ANNINE-6plus

One week after virus injection, GCaMP6f-expressing PNs were targeted for voltage-sensitive dye (ANNINE-6plus) single-cell labelling by electroporation, guided by two-photon microscopy (Kuhn and Roome, 2019). GCaMP6f-expressing PN were electroporated using a patch pipette containing 3 mM ANNINE-6plus dissolved in ethanol. Borosilicate glass (patch) pipettes with 1 μm tip diameter (7–10 MΩ) were used for electroporation and a stimulus protocol of 50 negative current pulses (−30 μA), 1 ms in duration at 100 Hz were delivered. Neutral pressure was applied to the patch pipette to prevent leakage of the dye/ethanol solution into tissue and the pipette was retracted immediately after the cell was loaded and replaced for further single-neuron labelling.

Typically, two to five PNs were filled per mouse on the same day, and after loading PNs with dye, mice were returned to their cages to allow the dye to spread to distal dendrites and throughout the entire cell. ANNINE-6plus is highly lipophilic so dye diffusion can take several hours (>12 h). After ~20 h, the brightest labelled cells were selected for imaging experiments, which were performed the day following PN labelling. This also guaranteed that the PN was healthy and had not been damaged by the labelling procedure. Where possible, several PNs were labelled in the same mouse (up to 5) and used for simultaneous dendritic voltage and calcium imaging recordings. Between neuron labelling sessions and following surgery, mice were returned to their cages and allowed to recover.

### Simultaneous dendritic voltage and calcium imaging

Approximately 20 h after ANNINE-6plus labelling, simultaneous imaging from PN dendrites was performed (<50 μm below dura) in linescan mode at 2 kHz sampling rate. Labelled PNs were clearly visible in both red and green channels indicating successful labelling with ANNINE-6plus and GCaMP6f. During recordings, mice were alert and headfixed sitting on a rotating treadmill. Mice were allowed to sit awake on the treadmill for at least 30 minutes before beginning the experiment. Bidirectional linescans, 512 pixels in width, lasting 10.5 s were performed at a line rate of 2 kHz. To limit photo-damage during linescans and to improve signal-to-noise ratio, the objective collar was rotated to elongate the excitation volume predominantly in the z-direction to ~5 μm. During the 10.5 s, no bleaching was observed. Fine corrections in linescan orientation (with respect to PN dendrites) were done prior to the experiment using the rotating stage, on which the mouse treadmill and micro-manipulators were placed. The linescans measuring 256 μm in width were carefully positioned as superficial as possible as to include the full dendritic width of the most distal dendritic spiny PN branches, thus maximizing the total membrane area covered by the linescan, typically less than 50 μm below the pia mater. ANNINE-6plus is purely electrochromic, showing linear responses across the full physiological voltage range and is well suited for recording neuronal membrane potential, with a temporal resolution limited only by the fluorescence lifetime (Fromherz et al., 2008). The femtosecond-pulsed Ti:sapphire laser was used to excite fluorescence at 1020 nm, near the red spectral edge of absorption. To confirm optimal ANNINE-6plus sensitivity near the red spectral edge of absorption and the mechanism of voltage sensitivity (Kuhn et al., 2004), different excitation wavelengths were tested (Roome and Kuhn, 2018). Excitation near the red spectral edge of absorption to optimize voltage sensitivity allows for long-term simultaneous voltage and calcium dendritic recordings at least 500 s per recording session at different dendritic depths. As ANNINE-6plus is relatively hydrophobic compared to other voltage-sensitive dyes for intracellular application, the labelling lasts for at least 2 weeks (Roome and Kuhn, 2018, 2019). Due to an extended excitation point spread function (~1 ×1× 5 μm^3^) used to increase the signal-to-noise ratio, the voltage signal is the average membrane potential in this volume encompassing spines and dendritic shafts.

### Pharmacology

A micropipette inserted through the chronic cranial window access port was used for pharmacological manipulation. In this case, the micropipette: a beveled quartz micropipette (0.7 mm ID, Sutter Instruments) was placed ~50 μm below the PN soma for drug injections. Lidocaine (2%) (Sigma) and CNQX disodium salt (100 μM) (Tocris) were used to block Na^+^ channels and AMPA receptors (excitatory synaptic input) respectively. CGP 35348 (100 μM) (Tocris) was used to block GABA_B_ receptors. GABA_B_ receptors are predominantly localized to the imaging region (PF-PN synapses in the spiny dendrites) (Kulik et al., 2002), whereas GABAA receptors are expressed abundantly throughout the cerebellum, notably at basket cell terminals on the PN soma (Fritschy and Panzanelli, 2006). Blocking GABAA receptors with SR 95531 hydrobromide (100 μM) (Tocris) led to over-excitability of the PN dendrites and irregular spontaneous spiking behavior. AIDA (100 μM) (Tocris) was used to block mGluR1 activity. All drugs were dissolved in saline and applied by pressure injection (<0.5 psi) for 10 min prior to imaging and reduced to <0.1 psi while dendritic voltage and calcium recordings were repeated in the awake mice. Dendritic recordings were occasionally repeated ~24h after drug application (guaranteeing drug washout) to confirm that the labelled PN was not physically damaged by the drug application.

### Data analysis

Linescan TIFF images were initially cropped in width using ImageJ (US National Institute of Health) to contain only the dendrite from the labelled PN using the red channel as a guide, and to eliminate green signals originating from neighboring PNs. Full linescan traces were imported into Matlab and interpolated (from 2 to 10 kHz). All subsequent data analysis was performed using custom-written programs that we wrote previously with Matlab (MathWorks). Minor movement corrections were made to linescan images in the spatial dimension, to minimize movement artefacts that occur during the sensory stimulus. This was achieved by maximizing the cross correlation between the average pre-stimulus linescan spatial profile, and the spatial profile of individual lines collected during the stimulus.

Single unit electrophysiological recordings were analyzed using Spike2 spike detection software (CED) to create a binary trace for simple spikes (SS) (and to eliminate spikes originating from neighboring PNs). All complex spikes (occurring at ~1 Hz) were easily identified by eye, which could be confirmed by comparing simultaneous electrophysiology, voltage and calcium traces. Complex spike binary traces were created using the initial sodium spike of the somatic complex spike to mark its onset. All binary traces, raw images and electrophysiology were then imported into Matlab for analysis.

To calculate ΔF/F of full linescans, the green channel image was first scaled (using Matlab ‘regress’ function) and fit to the red channel and then subtracted from the red channel. This removed crosstalk from the green channel. Average baseline red channel fluorescence was then subtracted from the time-varying red channel fluorescence and the result was divided by the average baseline of red channel fluorescence to give ΔF/F for the red channel. The ΔF/F calculation for the green channel was made in the same way as for the red channel, but it was not necessary to first subtract red channel fluorescence from the green channel as the voltage signal was neglectable compared to GCaMP6f. Relative fluorescence changes imaged with an excitation wavelength of 1020 nm were converted with a factor of 2.1 mV/% to estimate voltage changes.

### DCS and DS spikelet detection in Matlab

DCS and single DS signals were detected using a custom-written program in Matlab. Spatially averaged voltage and calcium traces were first temporally filtered. Voltage traces were filtered using a 1ms boxcar filter, while calcium traces were filtered using a 10ms boxcar filter. All spikelets were then detected in the voltage traces using the ‘findpeaks’ function in Matlab, for which ‘MinPeakHeight’ was set at 3 standard deviations of the voltage recording, the ‘MinPeakDistance’ was set at 2.5ms and ‘MinPeakProminence’ was set at 1 standard deviation of the voltage recording. Spikelets were then sorted into DS or DCS signals based on the following criteria:

Spikelets were classed as DS signals if:

1. No additional spikelets were detected in a window of ±10ms prior to and following the spikelet (i.e. no burst of spikelets was detected).
2. The average corresponding calcium signal measured in a 20ms window following the spikelet increased by > 1%, compared to the average calcium signal measured in a 20ms window prior to the spikelet (i.e. corresponding calcium signal was detected).
3. The average voltage signal measured in a 10ms window following the spikelet was less than 1 standard deviation of the voltage recording (i.e. the voltage recording returned immediately to baseline after a single spikelet).

Spikelets were classed as DCS signals if:

1. Additional spikelets were detected in a window of ±10ms prior to and following the spikelet (i.e. a burst of two or more spikelets were detected).
2. The average corresponding calcium signal measured in a 20ms window following the spikelet increased by > 1% ΔF/F, compared to the average calcium signal measured in a 20ms window prior to the spikelet (i.e. corresponding calcium signal was detected).

Classifications of DCS and DS signals were subsequently cross checked by visual inspection and though application of the classification algorithm, to paired dendritic and somatic recordings made on 7 PNs in awake mice (Roome and Kuhn, 2018), to confirm that DCS signals have a corresponding CF evoked somatic complex spike, while DS signals had no corresponding somatic signal (i.e. only dendritic) (see Figure 1 – figure supplement 2).

### Behavior analysis

Behavioral recordings were made while the mouse was awake and sitting on a treadmill. A behavior camera (sCMOS), capturing frames at 100 Hz, was used to record sensory stimulus evoked movements, synchronized with the imaging. Linescan recordings (10.5 seconds in length) included a 100ms sensory stimulus beginning after 5 seconds of recording, giving an air puff directed at the ipsilateral eye of the mouse. Air-puffs (pressure: 30psi) were delivered via a (0.7mm I.D) glass capillary positioned 2 cm from the eye to evoke a reliable eye blink reflex in the ipsilateral eye. Regions of interest (ROIs) were drawn around the ipsilateral eye and the Δintensity/intensity was used to quantify eyelid position and closure during the stimulus, and traces were normalized such that 1 and 0 correspond to the maximum and minimum eyelid closure recorded for each mouse across all recordings, in order to monitor changes in the eye response over time.

## Acknowledgments

We thank the GENIE Program and the Janelia Research Campus for distributing GCaMP6f, Lina Koronfel and Soumen Jana for helpful feedback on the manuscript. We are grateful for generous support and funding from the Okinawa Institute of Science and Technology Graduate University.

## Author contributions

C.J.R. designed the study, built the setup, collected and analyzed data, and wrote the paper. B.K. designed the study, built the setup, and wrote the paper.

## Figure supplements

**Figure 1 – figure supplement 1.**
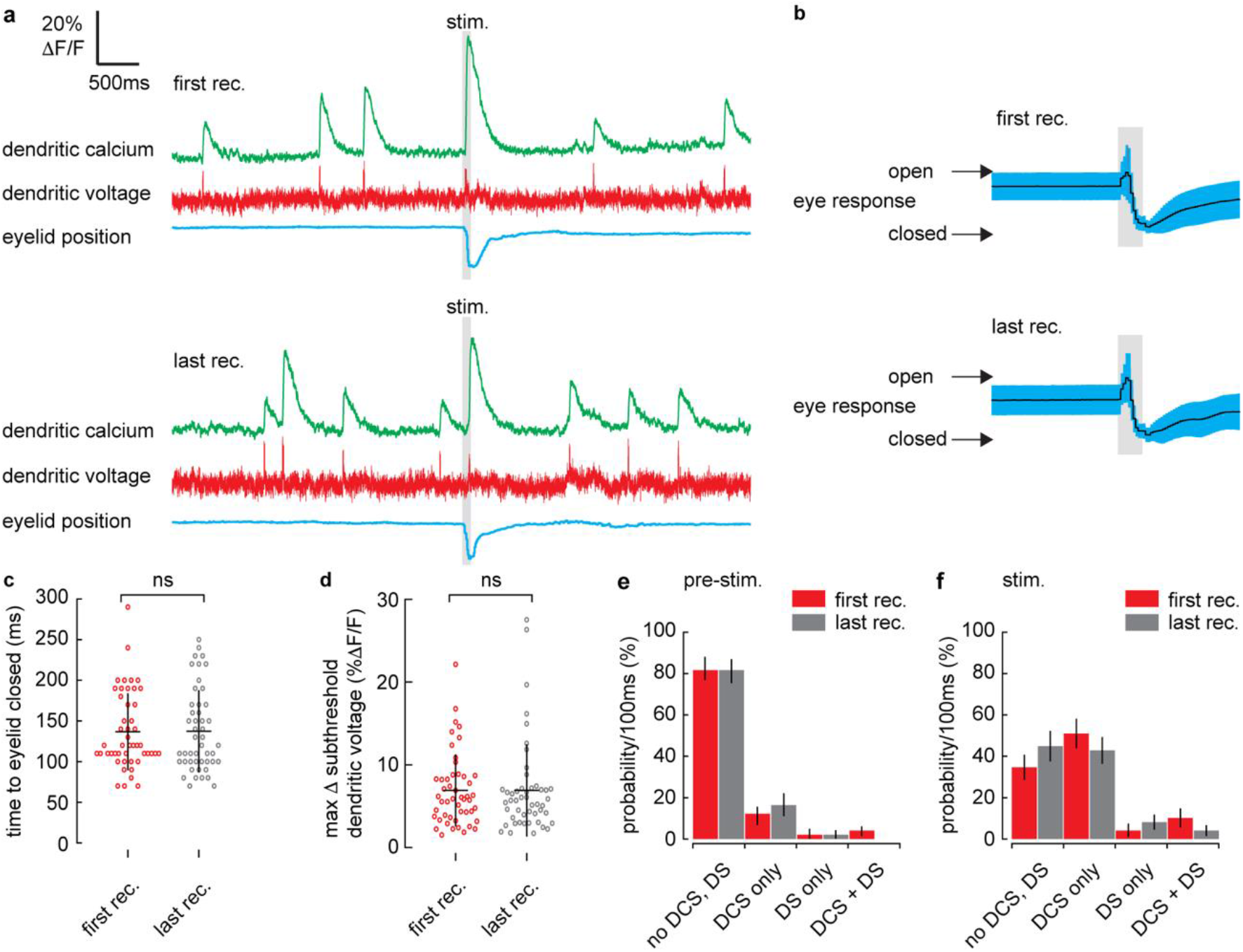
No difference in subthreshold or suprathreshold voltage signaling or time to eyelid closed between first and last recordings. (a) Single trial recordings of simultaneous dendritic voltage (red traces) and calcium (green traces) during sensory stimulation (stim.). Suprathreshold signals are clearly visible in both voltage and calcium traces. Blue traces show average intensity of ROI used to record eyelid position (eyeblink responses). Grey bars show full duration of air puff stimulus. Top example shows voltage and calcium during first recording (rec.) and bottom example shows last (10^th^) recording from the same PN. (b) Mean eye responses from first (top; n = 49) and last (bottom; n = 49) recordings (black trace shows mean; blue trace shows SD). (c) Time for eyelid to close after stimulus onset for first (red) and last (grey) recording groups. There was no significant difference in eyelid closure time between first (136 ± 47 ms, mean ± SD), and last (138 ± 50 ms, mean ± SD) groups, p = 0.97, Wilcoxon signed rank test, 49 pairs from 49 PNs. (d) Maximum change in subthreshold voltage measured during the stimulus from first and last recordings. Stimulus evoked subthreshold voltage signals from first (6.8 ± 4.3 % ΔF/F, mean ± SD), and last (6.6 ± 5.3 mean ± SD) recording groups were not significantly different, p = 0.886, unpaired t-test, 49 pairs from 49 PNs. Probability distributions for suprathreshold DCS and DS signals during pre-stimulus (d) and stimulus (e) time windows, sorted into first and last recording groups (as in (c) and (d)). There were no significant differences in probability distributions between first and last recording groups, during both pre-stimulus and stimulus time windows, Kolmogorov-Smirnov, p > 0.95, 49 pairs from 49 PNs. Vertical bars in probability distributions show bootstrapped SD.

**Figure 1 – figure supplement 2.**
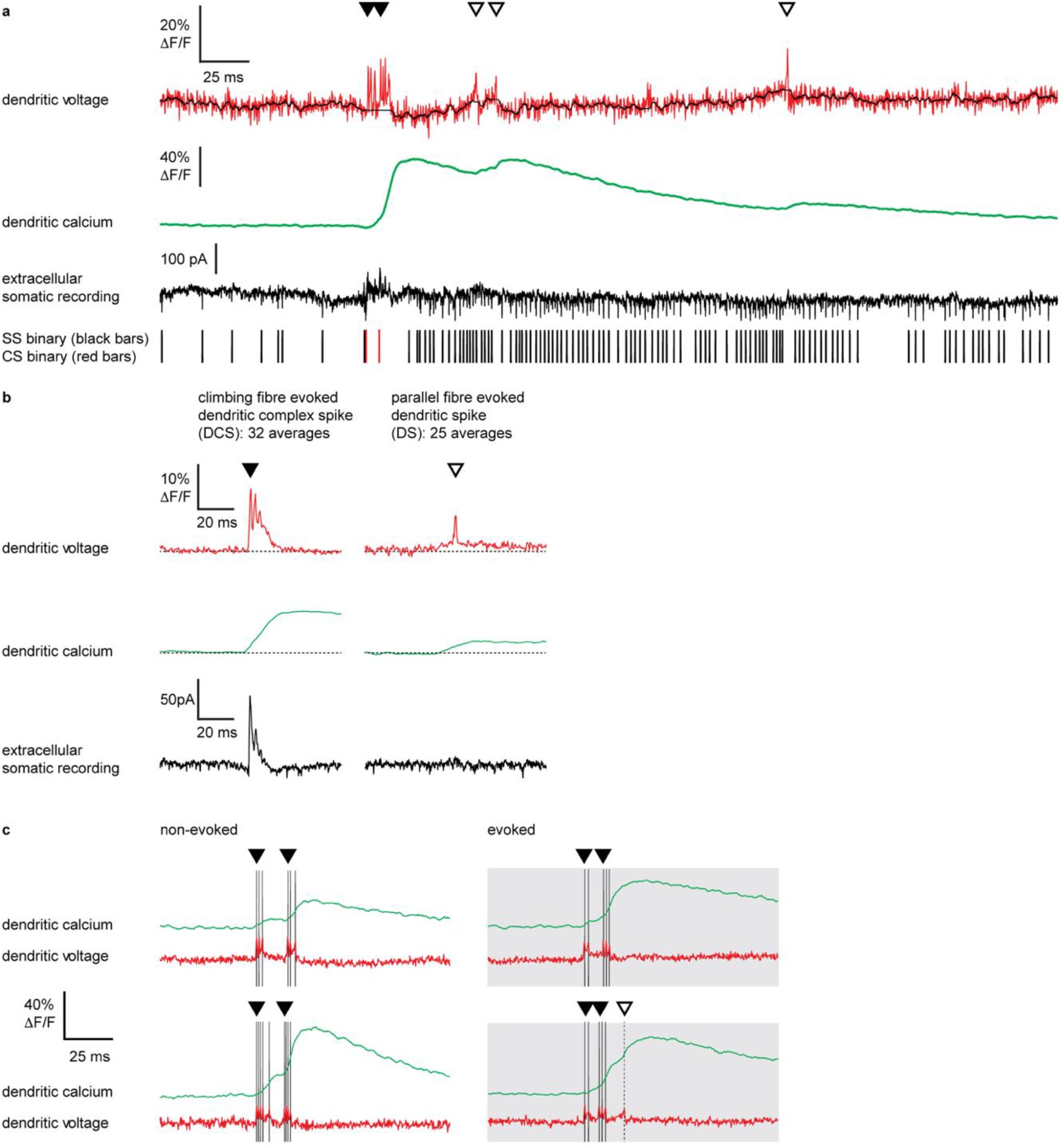
Suprathreshold DCS and DS detection in Purkinje neuron dendrites of awake mice. (a) Single trial recordings of simultaneous dendritic voltage (red trace, black overlay shows subthreshold voltage, with spikelets removed and 5ms boxcar smoothing) and calcium (green trace) and paired extracellular somatic recording (black trace). Binary traces for somatic simple spikes (SS; black bars) and complex spikes (CS; red bars) are shown. (b) averages of 32 DCS signals (left) and 25 DS signals (right), showing dendritic voltage (red), dendritic calcium (green) and extracellular somatic recording (black). Note that DCS signals are characterized by a burst of dendritic spikelets, a large calcium transient, and corresponding complex spike at the soma, while DS signals are characterized by a gradual membrane depolarization, a single spikelet, and small calcium transient in the dendrites, and no corresponding signal at the soma. (c) Single trial recording examples of non-evoked (left) and evoked (right) dendritic voltage spike signals (red) and corresponding dendritic calcium signals (green) during high frequency DCS and DS activity. Black and dashed bars show detected spikelets during DCS and DS signals respectively. DCS signals are indicated by filled black triangles and DS signals are indicated by open triangles.

**Figure 1 – figure supplement 3.**
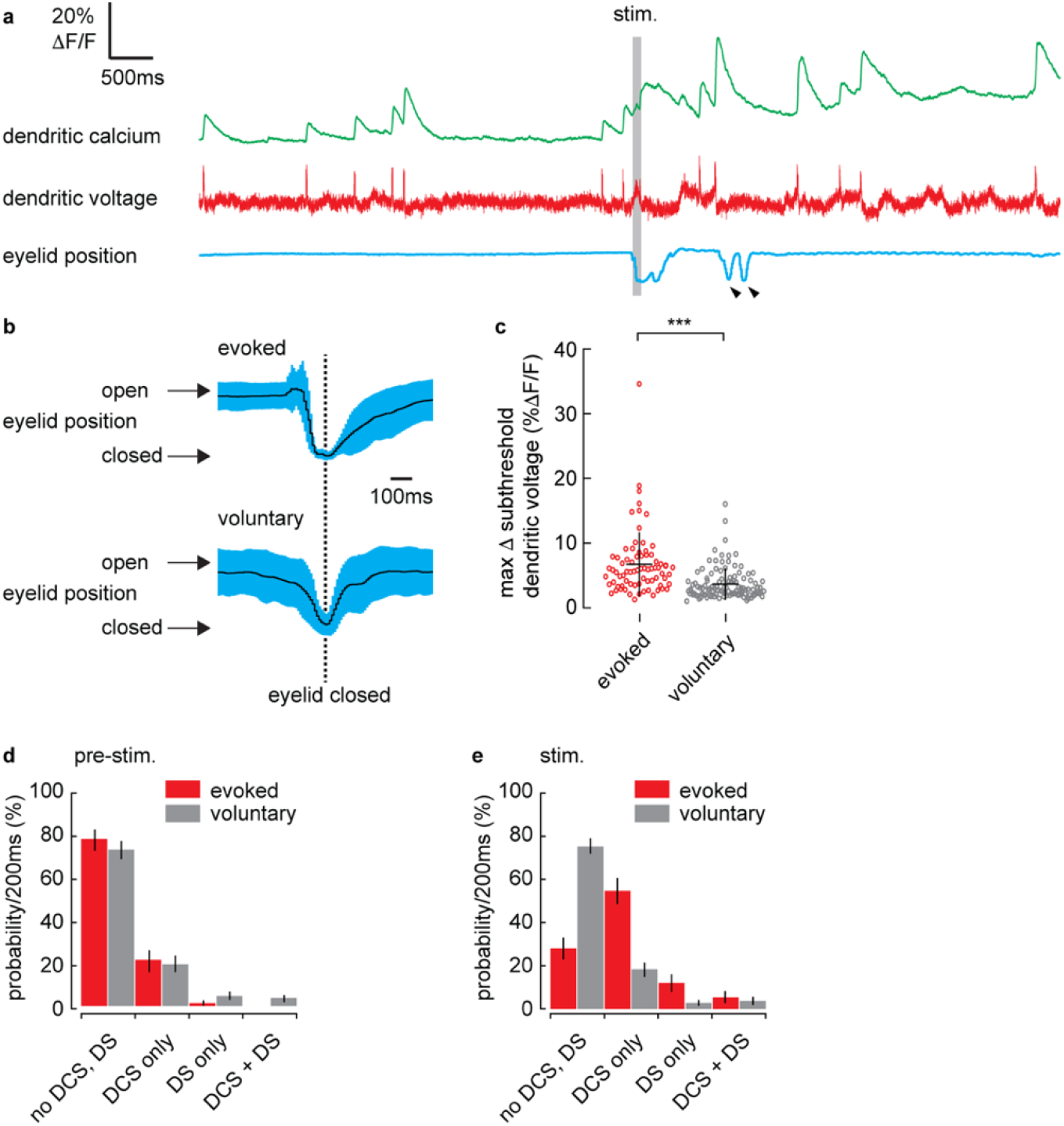
Subthreshold and suprathreshold dendritic voltage signals are time locked to sensory stimuli and not voluntary eye blinks. (a) Single trial recordings of simultaneous dendritic voltage (red trace) and calcium (green trace) during sensory stimulation (stim.). Suprathreshold signals are clearly visible in both voltage and calcium traces. Blue traces show average intensity of ROI used to record eyelid position (eyeblink responses). Grey bar shows full duration of air puff stimulus. Small black arrows indicate ‘voluntary’ eye blinks occurring > 1s after stimulus ends. For comparison only the recordings containing both evoked and voluntary eye blinks were selected, (n = 75, 24 PNs). (b) Mean eye responses from evoked (top) and voluntary (bottom) eye blinks aligned to full eyelid closure (dashed black line) (black trace shows mean; blue trace shows SD). (c) Maximum change in subthreshold voltage in a 200ms window preceding full eyelid closure during evoked and voluntary eye blinks. Stimulus evoked subthreshold voltage signals were significantly larger during evoked eyeblinks; control (6.7 ± 5.0 % ΔF/F, mean ± SD, n = 75), compared to voluntary eyeblinks (3.7 ± 2.4 % ΔF/F, mean ± SD, n = 109), unpaired t-test, p = 1.1×10^−7^, n = 75, 24 PNs. Probability distributions for suprathreshold DCS and DS signals during a pre-stimulus (d) and stimulus (e) time window, for evoked and voluntary eyeblinks. Pre-stimulus time windows are defined here as a 200ms window, beginning 500ms before eye closure. Stimulus time windows are defined here as a 200ms window immediately preceding eyelid closure. There was no difference between evoked and voluntary eyeblink probability distributions during pre-stimulus time windows (pre-stimulus, p = 0.98). During stimulus time windows the evoked eyeblink probability distribution was shifted towards a higher probability of suprathreshold signals compared to voluntary eyeblinks (stimulus, p = 2.17×10^−9^). Kolmogorov-Smirnov, n = 75, 24 PNs. Vertical bars in probability distributions show bootstrapped SD. *** indicate p < 0.001.

**Figure 1 – figure supplement 4.**
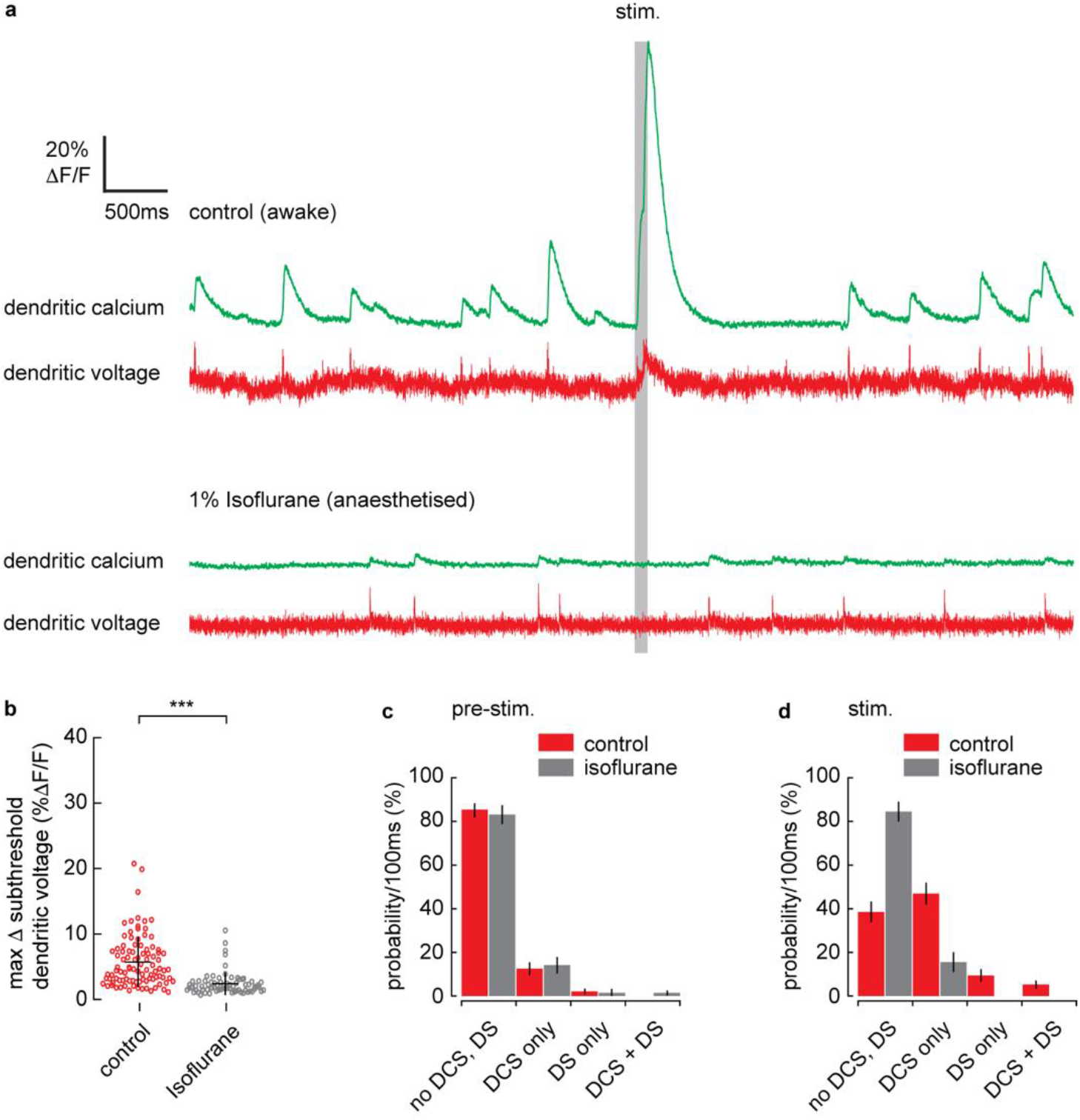
Isoflurane induced anesthesia blocks stimulus evoked subthreshold and suprathreshold dendritic voltage signaling. (a) Single trial recordings of simultaneous dendritic voltage (red traces) and calcium (green traces) during sensory stimulation (stim.). Suprathreshold signals are clearly visible in both voltage and calcium traces. Grey bars show full duration of air puff stimulus. Top example shows voltage and calcium recordings during control (awake) conditions and bottom example shows recordings during anesthesia (1% Isoflurane) from the same PN. (b) Maximum change in subthreshold voltage measured within a 200ms window after stimulus onset under control (red) and Isoflurane (grey) conditions. Stimulus evoked subthreshold voltage signals were significantly reduced during anesthesia; control (5.7 ± 3.9 % ΔF/F, mean ± SD, n = 96), and Isoflurane (2.4 ± 1.7 % ΔF/F, mean ± SD, n = 71), unpaired t-test, p = 2.4×10^−10^, n = 167, 6 PNs. Probability distributions for suprathreshold DCS and DS signals during (c) pre-stimulus and (d) stimulus time windows, under control and Isoflurane conditions. Probability distributions during pre-stimulus time windows were unaltered by isoflurane; (pre-stimulus, p = 1.0). Probability distributions during stimulus time windows were shifted towards a higher probability of no-signals during isoflurane anesthesia, (stimulus; p = 3.1×10^−8^). Unlike in awake conditions, during anesthesia there was no difference between pre-stimulus and stimulus probability distributions (p = 1.0). Kolmogorov-Smirnov, n = 167, 6 PNs. Vertical bars in probability distributions show bootstrapped SD. *** indicate p < 0.001.

**Figure 1 – figure supplement 5.**
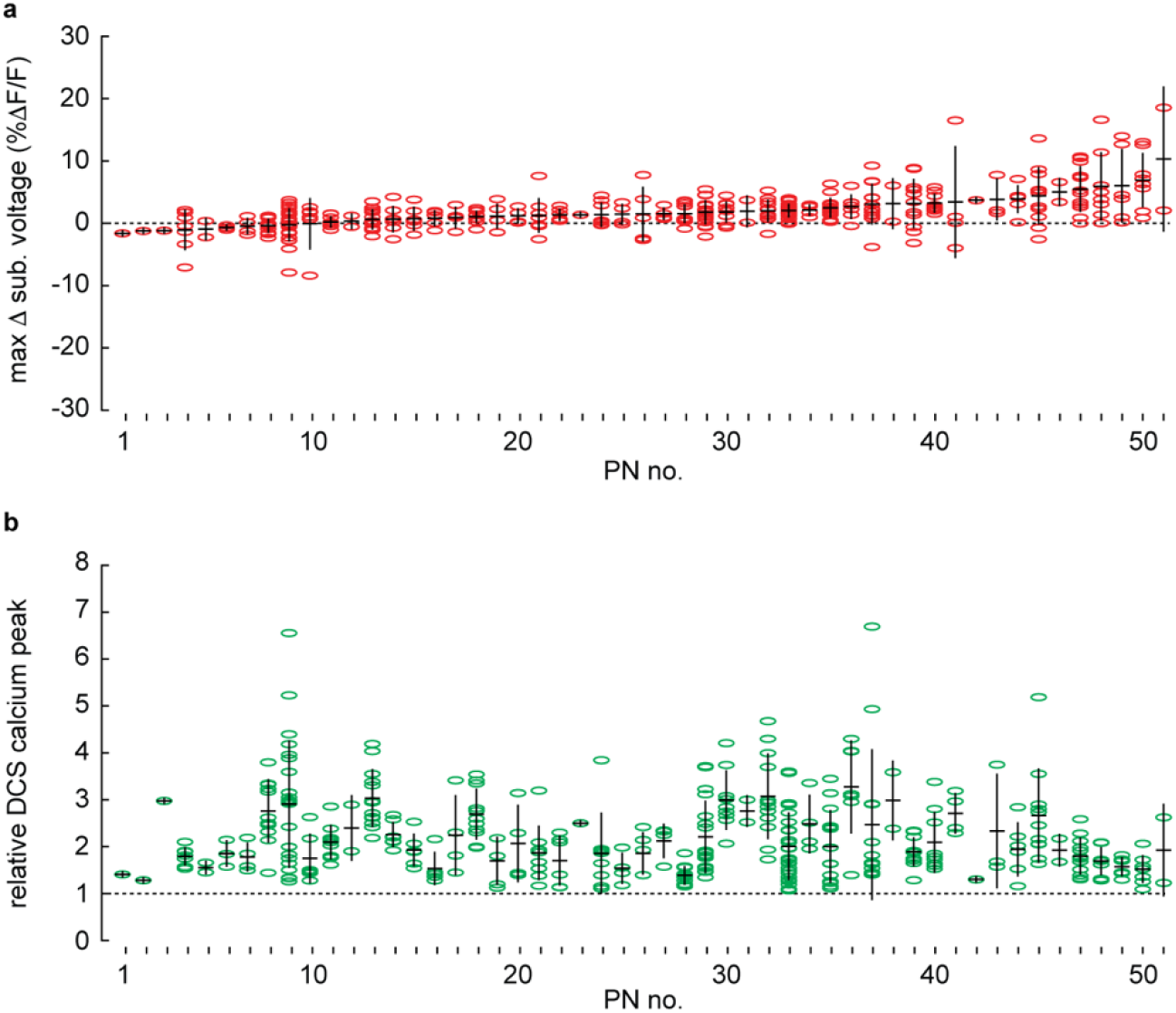
During sensory stimulation the average maximum change in subthreshold voltage is positive and dendritic calcium signals are enhanced. (a) Maximum change in subthreshold voltage during sensory stimulation measured from all recordings with evoked DCS signals, n = 365, 51 PNs (sorted in ascending order based on average for each PN); group average; 2.0 ± 2.3 % ΔF/F, is significantly above baseline (dashed line), p = 2.0×10^−7^, t-test. (b) Stimulus evoked DCS associated calcium signals, relative to the average non-evoked DCS calcium signal for each PN, n = 365, 51 PNs, using the same sorting as in (a), group average; 1.4 ± 0.6, is significantly above baseline (dashed line), p = 1.5×10^−5^, t-test.

**Figure 3 – figure supplement 1.**
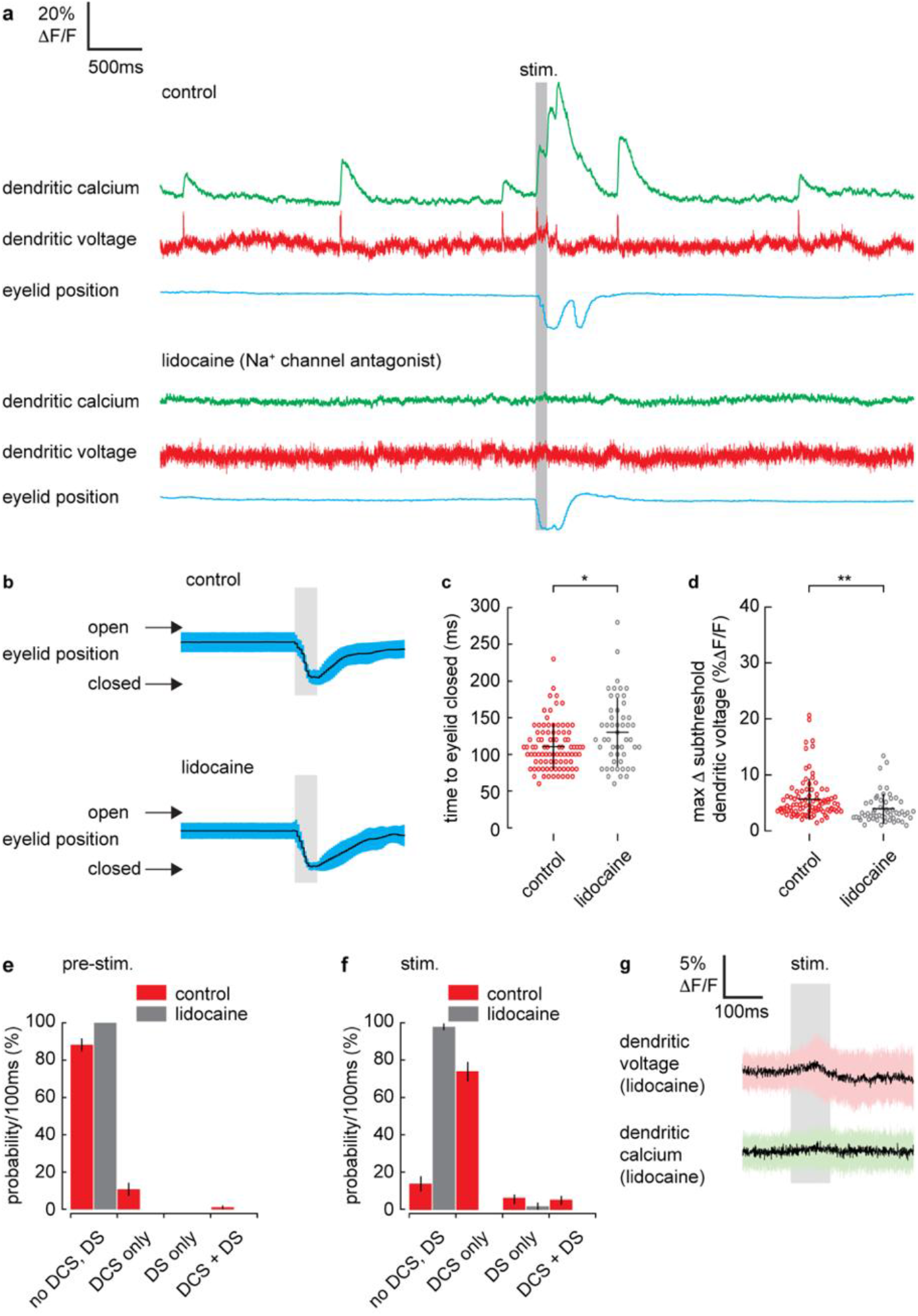
Na^+^ channel antagonist, Lidocaine, blocks stimulus evoked subthreshold and suprathreshold dendritic voltage signaling and slows stimulus evoked eye blinks. (a) Single trial recordings of simultaneous dendritic voltage (red traces) and calcium (green traces) during sensory stimulation (stim.). Suprathreshold signals are clearly visible in both voltage and calcium traces. Blue traces show average intensity of ROI used to record eyelid position (eyeblink responses). Grey bars show full duration of air puff stimulus. Top example shows voltage and calcium during control conditions and bottom example shows recordings following Lidocaine application from the same PN. (b) Mean eye responses from control (top, n = 93) and Lidocaine (bottom, n = 54) recordings (black trace shows mean; blue trace shows SD). (c) Time for eyelid to close after stimulus onset under control (red) and Lidocaine (grey) conditions. There was a small but significant increase in eyelid closure time following Lidocaine application; control (111 ± 31 ms, mean ± SD, n = 90); Lidocaine (131 ± 46 ms, mean ± SD, n = 50), Kolmogorov-Smirnov, p = 0.02, n = 140, 6 PNs. (d) Maximum change in subthreshold voltage measured within a 200ms window after stimulus onset under control and Lidocaine conditions. Stimulus evoked subthreshold voltage signals were significantly reduced following Lidocaine application; control (5.7 ± 3.7 % ΔF/F, mean ± SD, n = 93), and Lidocaine (3.9 ± 2.7 % ΔF/F, mean ± SD, n = 54), unpaired t-test, p = 0.003, n = 147, 6 PNs. Probability distributions for suprathreshold DCS and DS signals during pre-stimulus (e) and stimulus (f) time windows, under control and Lidocaine conditions (as in (c) and (d)). Lidocaine blocks suprathreshold signals during both pre-stimulus and stimulus time windows. There were no suprathreshold signals detected during the pre-stimulus time window following lidocaine application and probability distributions were shifted towards a higher probability of no-signal following Lidocaine application during the stimulus time window, Kolmogorov-Smirnov, p = 1.8×10^−22^, n = 147, 6 PNs. Vertical bars in probability distributions show bootstrapped SD. * and ** indicate p < 0.05 and p < 0.01 respectively. (g) Averages of subthreshold voltage (red) and calcium (green) dendritic signaling during sensory stimulation, following Lidocaine application, black line shows mean, colored trace shows SD. Calcium at stimulation offset was not significantly different from baseline, t-test, p = 0.2, n = 51, 6 PNs.

**Figure 7 – figure supplement 1.**
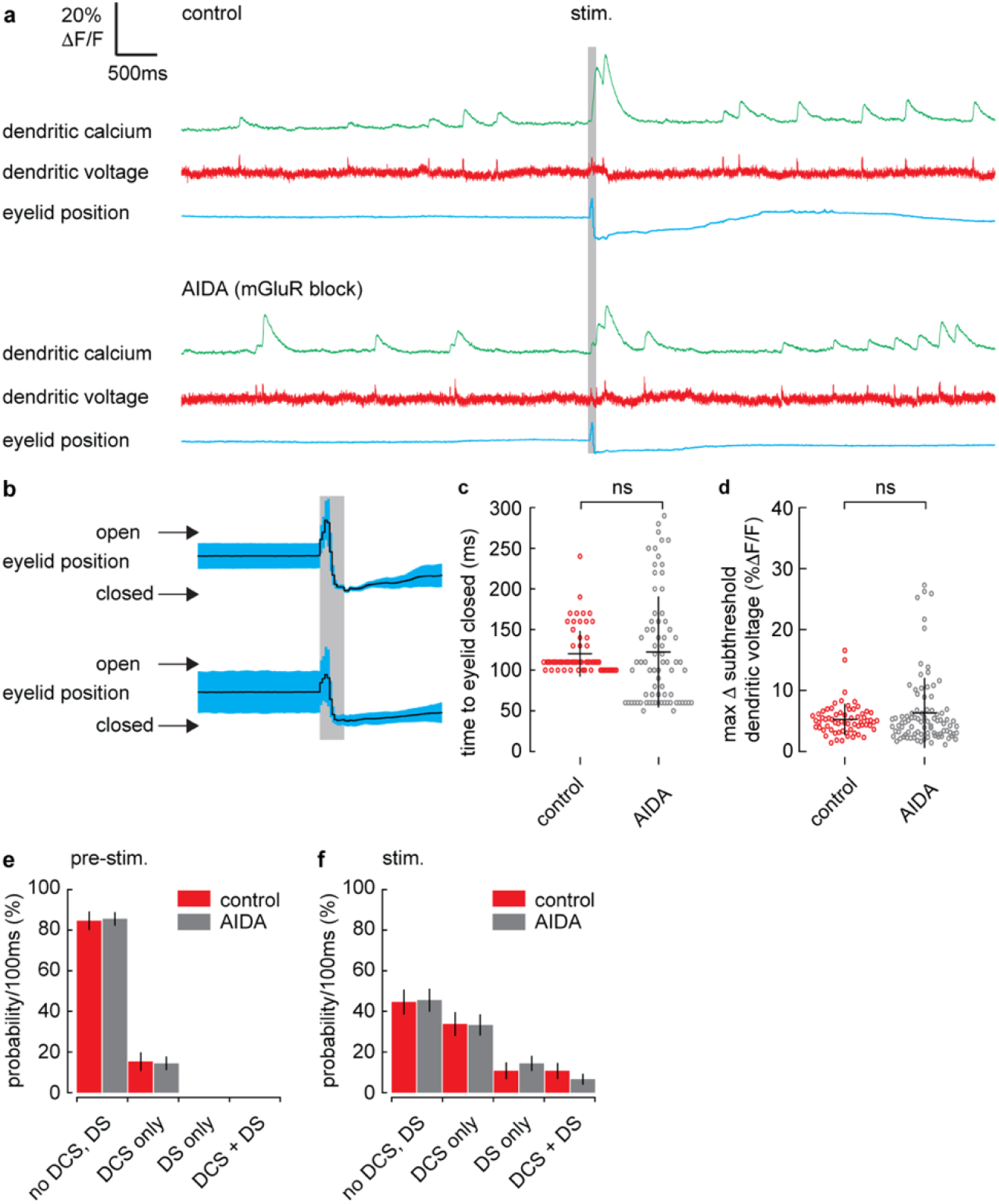
mGluR antagonist, AIDA, has no effect of evoked eye blink closure time, stimulus evoked subthreshold voltage signaling or the probability of suprathreshold dendritic signals. (a) Single trial recordings of simultaneous dendritic voltage (red traces) and calcium (green traces) during sensory stimulation (stim.). Suprathreshold signals are clearly visible in both voltage and calcium traces. Blue traces show average intensity of ROI used to record eyelid position (eyeblink responses). Grey bars show full duration of air puff stimulus. Top example shows voltage and calcium during control conditions and bottom example shows recordings following AIDA application from the same PN. (b) Mean eye responses from control (top) and Lidocaine (bottom) recordings (black trace shows mean; blue trace shows SD). (c) Time for eyelid to close after stimulus onset under control (red) and AIDA (grey) conditions. There was no significant difference in eyelid closure time following AIDA application; control (120 ± 28 ms, mean ± SD, n = 64); AIDA (122 ± 68 ms, mean ± SD, n = 74), Kolmogorov-Smirnov, p = 0.29, n = 138, 6 PNs. (d) Maximum change in subthreshold voltage measured within a 200ms window after stimulus onset under control and AIDA conditions. Stimulus evoked subthreshold voltage signals did not change following AIDA application; control (5.3 ± 2.6 mean ± SD, n = 65), and AIDA (6.3 ± 5.8, mean ± SD, n = 90), unpaired t-test, p = 0.17, n = 155, 6 PNs. Probability distributions for suprathreshold DCS and DS signals during pre-stimulus (e) and stimulus (f) time windows, under control and AIDA conditions (as in (c and d)). AIDA did not alter the probability of suprathreshold signals during either pre-stimulus or stimulus time windows, (pre-stimulus, p = 1.0); (stimulus, p = 1.0), Kolmogorov-Smirnov, n = 155, 6 PNs. Vertical bars in probability distributions show bootstrapped SD.

